# Inferring tumour microenvironment ecosystems from scRNA-seq atlases

**DOI:** 10.1101/2024.08.21.608956

**Authors:** Chengxin Yu, Michael J Geuenich, Sabrina Ge, Gun-Ho Jang, Tan Tiak Ju, Amy Zhang, Grainne M O’Kane, Faiyaz Notta, Kieran R Campbell

## Abstract

Accurate inference of granular cell states that co-occur within the tumour microenvironment (TME) is central to defining pro– and anti-tumour environments. Here, we describe how ecosystems of cell populations can be robustly inferred from nonspatial single-cell RNA-seq (scRNA-seq) atlases. Leveraging a unique discovery-validation setup across eight scRNA-seq datasets profiling pancreatic ductal adenocarcinoma (PDAC), we show highly consistent co-occurrence of fine-grained cell states across patients and characterize the positive predictive value of such analyses. Building on this, we develop a novel probabilistic model to quantify multi-cellular ecosystems directly from such atlas-scale scRNA-seq datasets. By mapping these ecosystems to spatial transcriptomics data, we demonstrate that such ecosystems represent bona fide spatial variation within individual tumours despite being learned from nonspatial data. Importantly, through mapping these ecosystems across two large clinical cohorts, we show they are more predictive of therapy response than any individual cell state and are associated with specific tumour somatic mutations. Together, this work lays the foundation for inferring reproducible multicellular ecosystems directly from large nonspatial scRNA-seq atlases.

## Introduction

Recent advances in expression profiling technologies have significantly accelerated our ability to quantify cellular phenotypes at single-cell and spatial resolutions. Single cell RNA-seq^1,2^ (scRNA-seq) enables transcriptome-wide gene expression measurements at single cell resolution and has led to multiple discoveries of cell types and transcriptional states associated with patient outcomes and therapy response^3–8^. In tandem, spatial expression profiling technologies^9–12^ that quantify gene or protein expression at (sub-) cellular resolution while retaining the spatial location of each measurement have led to multiple studies that associate tumour spatial architecture with patient outcomes^4,5,13,14^. Of note, significant attention has been given to using these technologies for the *de novo* identification of cell types and states that co-occur within tissues^15^ and form multi-cellular communities (or ecosystems)^16–18^. These ecosystems form multi-cellular networks whose presence in tissues has been linked to clinical outcomes such as treatment response across a range of pathologies^13,19^. Importantly, much of our understanding of tissue dynamics leading to target discovery comes from reasoning about cellular co-occurrence within the TME.

However, reliable quantification of cell state co-occurrence within individual tumours leading to cellular ecosystems remains challenging. In particular, the ability to robustly identify recurrent coexisting cell states or gene programs in specific clinical scenarios relies on having sufficient independent samples, particularly for rare or transient cell states. Spatial protein technologies are sufficiently cost-efficient to profile the required number of tissue samples, but can only quantify 40-60 markers simultaneously, limiting the ability to infer granular cell states and signaling pathways. On the other hand, spatial transcriptomic technologies can measure 100s of mRNA species or even the whole transcriptome simultaneously, but are currently too expensive to profile at scale and suffers resolution or RNA contamination problems^20^. In contrast, scRNA-seq can quantify single-cell whole transcriptome expression measurements with quantified cell state abundance largely reflecting tissue composition^21–23^. While scRNA-seq nominally suffers from the same expense per sample as spatial transcriptomics technologies, the push by the community over the past decade through initiatives such as the Human Cell Atlas^24^ and Human Tumour Analysis Network^25^ has resulted in unprecedented data sizes, comprising more than 12 tumour atlases^4,26–31^ by mid-2024 alone. This leads to the tantalizing possibility of quantifying cell state co-occurrence and cellular ecosystems directly from large multi-cohort scRNA-seq cancer atlases. However, whether this can lead to reproducible, clinically relevant insights remains largely unknown.

To help answer this question, we focus on pancreatic ductal adenocarcinoma (PDAC) due to both the availability of multiple independent primary tumour scRNA-seq datasets studies^4,32–38^, combined with its complex microenvironment that is characteristically difficult to target. Existing studies of the PDAC TME have highlighted diverse cellular heterogeneity across all lineages^39– 43^. Additionally, single-cell profiling of PDAC tumours has highlighted previous underappreciated heterogeneity within individual tumours cells themselves, with the basal-like/classical^44^ and later hybrid subtypes^40^ being shown to co-exist within single tumours^45,46^. Despite this, few efforts have sought to explicitly quantify the co-occurrence of granular cell states within single tumours. Oh et al. curated scRNA-seq PDAC from four studies to infer cell type co-occurrence using clustering and interactome analysis^41^, but due to having relatively few samples did not include validation or highly granular cell states. In contrast, Luca et al. built a machine learning framework EcoTyper to discover multicellular communities across solid tumours from bulk RNA-seq data, which is well-powered at the patient level but may lack the ability to uncover granular cell states *de novo*. Therefore, there is an outstanding question for whether granular PDAC cell states can be robustly inferred and validated across patient cohorts, and whether cell state co-occurrence provides insights into the immune dynamics in tumours.

In this work, we present a comprehensive platform to dissect the extent to which cell state co-occurrence and multi-cellular ecosystems can be reproducibly inferred in the PDAC tumour microenvironment from atlas-scale scRNA-seq (**Fig. 1A**). We collate eight independent datasets encompassing >150 samples and >500,000 cells to create a unique dataset-level discovery-validation setup. In doing so, the existence of known and *de novo* granular cellular subtypes in the TME can be independently verified, along with their ability to co-occur within and across cell types and form cellular ecosystems. We map 50 cell type-specific gene programs across 8 cell types and confirm the existence of known subtypes such as exhausted cytotoxic T cell while discovering novel cellular variation such as multiple reproducible inflammatory cancer associated fibroblast (CAF) subtypes. Next, we evaluate the extent to which co-occurrence of these granular cell state programs is reproducible within and between cell types, finding consistent co-occurrence across major cell types. We subsequently formalize a probabilistic model for quantifying multicellular ecosystems from atlas-scale scRNA-seq and find them to be reproducible across our datasets. Interestingly, by mapping these ecosystems to spatial transcriptomics data, we demonstrate that such ecosystems represent bona fide spatial variation within individual tumours despite being learned from nonspatial data. Finally, we map the validated gene programs and ecosystems to two bulk datasets and find that certain ecosystems are associated with patient outcome along with specific KRAS mutations.

**Figure 1:**
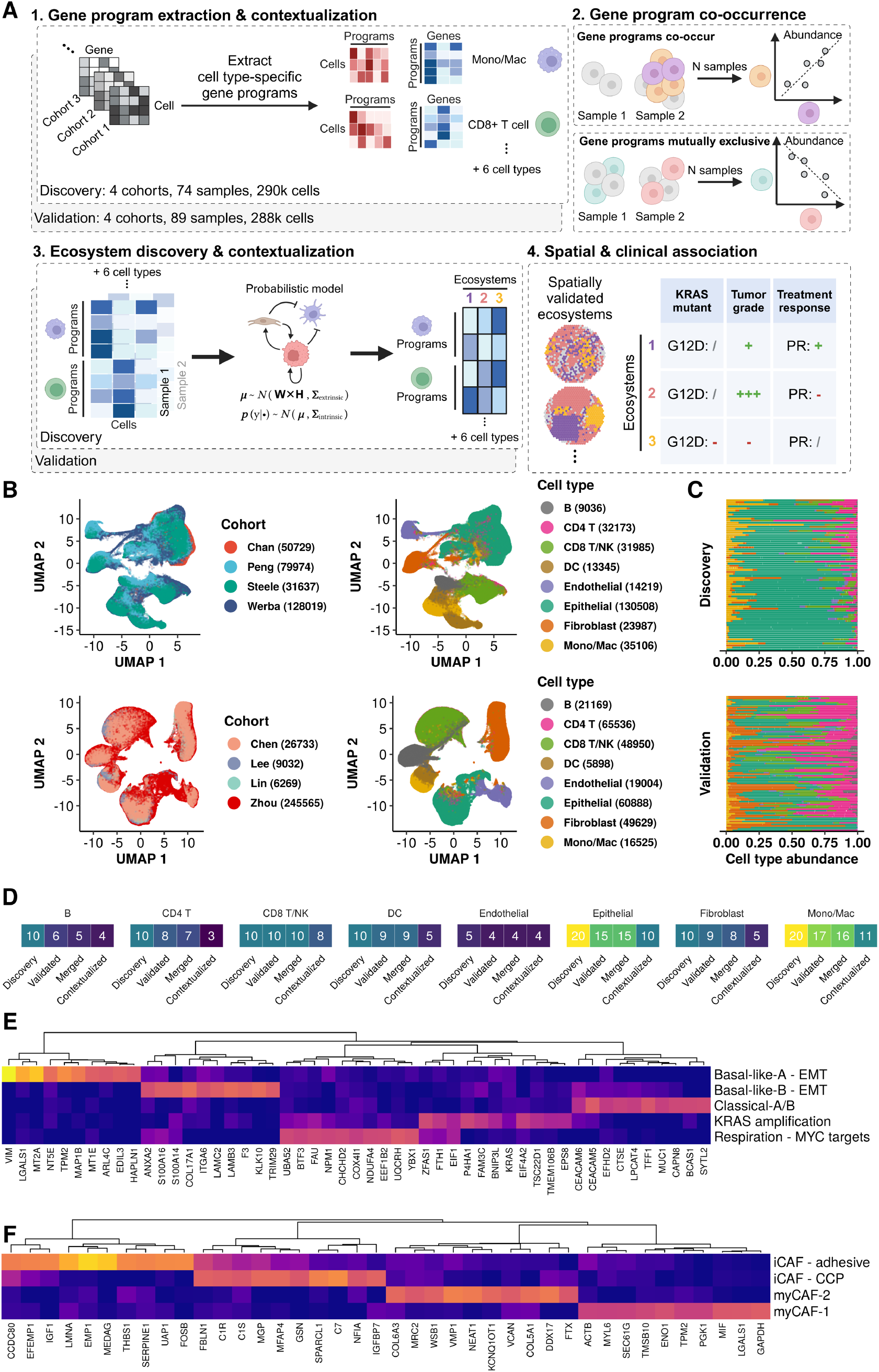
Learning multicellular ecosystems directly from scRNA-seq atlases. **A**. Overview of discovery-validation platform for identifying co-occurrence and ecosystems of cell type specific gene programs in the PDAC TME. **B**. UMAP representations for the discovery (top) and validation (bottom) datasets colored by cohorts (left) and annotated cell type (right). DC, Myeloid Dendritic Cells. Mono/Mac, Monocyte/Macrophage. **C**. Overview of cell type composition for samples (in rows) in discovery group (top) and in validation group (bottom). **D**, Overview of numbers of candidate gene programs extracted from the discovery group, numbers of gene programs validated between discovery and validation, and numbers of consolidated gene programs after merging the redundant ones in each cell type. **E-F**. Heatmaps showing weights of top ranked genes in selected gene programs from epithelial cells (**E**) and fibroblasts (**F**).

## Results

### Identifying reproducible cellular subtype heterogeneity in the PDAC TME

To systematically quantify cell states from the PDAC tumour microenvironment, we compiled 8 previously generated scRNA-seq datasets (**Supplementary Fig. 1A** and **Methods, Supplementary Table 1**)^4,32–37,47^ comprising 578,000 total cells. To exclude possible technology-induced bias, all selected scRNA-seq data were generated using the 10x Genomics platform. We selected primary PDAC tumour samples from each patient cohort and split the datasets into a discovery group (4 datasets, n=74 tumour samples) and a validation group (4 datasets, n=89 tumour samples). Raw count data preprocessing was performed in line with previous studies (**Supplementary Fig. 1B, Methods**). An automated workflow was developed to systematically annotate single cells using a public multi-cohort PDAC atlas as reference (**Supplementary Fig. 2A** and **Methods**)^47^. Eight major cell types were identified in our curated atlas (**Fig. 1B**). To validate the automatically assigned cell type labels, we examined known marker expressions in each dataset (**Supplementary Fig. 2B**).

To quantify fine-grained cellular subtypes in the TME, we extracted cell type-specific gene programs independently for the discovery and the validation datasets. Specifically, we applied integrative non-negative matrix factorization (iNMF)^48^ that learns a set of gene programs defined by the expression pattern of a set of genes along with their activities in each single cell (**Supplementary Fig. 3A, Methods**). The inherent plasticity of the cells of the TME motivated our use of NMF rather than sub-clustering^5,32,49,50^, which allows each cell to have a continuous activation of multiple gene programs compared to clustering approaches that force each cell to belong to a distinct cluster, following recent work in this area^51^.

Given concerns around whether granular cell subclusters from scRNA-seq reflect noise or overclustering^52^, we set out to identify a subset of cell type-specific high-confidence gene programs found in both our discovery and validation datasets. We first employed the Hungarian algorithm^53^ to obtain optimal one-to-one mapping between gene programs from the discovery and the validation datasets (**Supplementary Fig. 3B, C**). Next, we merged mapped gene programs with similar gene weights to reduce redundancy (**Methods: Program validation, Program merge**) (**Supplementary Figure 3D**). Lastly, we removed any gene programs likely corresponding to ambient RNA programs contaminating^54^ by manual examination of the top program markers (**Methods**). After applying this procedure, we obtained 10 high-confidence gene programs in epithelial cells, 5 high-confidence gene programs in each of fibroblasts and dendritic cells, 11 in monocytes/macrophages, 8 in CD8 T/NK cells, 3 in CD4 T cells, 4 in B cells, and 4 in endothelial cells (**Fig. 1D**). In total, we obtained 50 cell type-specific gene programs that capture both the overall TME landscape and the granular phenotypic heterogeneity (**Supplementary Fig. 4**). Overall, our approach allowed us to identify high-confidence recurrent cellular subtypes in PDAC TMEs for downstream analysis.

We next contextualized the high-confidence cell type-specific gene programs to distinguish potentially novel PDAC TME cell states from known ones. We took a multi-faceted approach including marker gene inspection, GSEA, and literature review (**Methods**). Importantly, our unique discovery-validation setup that independently discovered reproducible programs in multiple datasets confers high confidence on their manifestation in patients. While this recapitulated many existing cell states as previously reported in single-cell profiling of PDAC^32,39,44,55^—including basal and classical epithelial subtypes as well as inflammatory and myofibroblastic cancer associated fibroblasts (CAFS)—we discovered certain high-confidence gene programs previously undescribed.

In the epithelial compartment, we discovered a previously undescribed single-cell KRAS amplification epithelial program marked by upregulation of *KRAS* and *BNIP3L*/*NIX—*previously shown to be a driver of glycolysis and disease progression in pancreatic cancer^56^ (**Fig. 1E**). In the fibroblast compartment, we identified two distinct inflammatory cancer associated fibroblast (iCAF) subsets and two myofibroblastic CAF (myCAF) subsets for the first time (**Fig. 1F**). The *classical pathway (CCP) iCAF* program showed strong upregulation of *C7* and the classical pathway genes *C1S* and *C1R*. The CCP-iCAF program was also enriched for a neurotropic gene set linked to tumour growth, stress adaptation, and metastasis in PDAC^5,57^ (**Supplementary Fig. 5A**). In contrast the *adhesive iCAF* program was highly enriched for an adhesion gene signature including strong upregulation of the gene *SERPINE1* that encodes for the protein plasminogen activator inhibitor-1 (*PAI-1*) (**Fig. 1F**). The *ACSL3*-*PAI-1* signaling axis supports PDAC progression, links to fibrosis, immunosuppression, and prognosis^58^. We found the CCP iCAF program and the adhesive iCAF program to be present in a mutually exclusive pattern across patients within the dataset (**Supplementary Fig. 5B**). Two myCAF programs were identified based on CAF subtype gene loadings in fibroblast gene programs (**Supplementary Fig. 5C**). The *TPM2+* myCAF-1 program was highlighted by the high loading of *MIF*^56^ (**Fig. 1F**). The *VMP1*+ myCAF-2 program was identified as a potential novel myCAF subtype. Although VMP1 has been associated with PanIN formation^59^ and patient survival in pancreatic neuroendocrine tumours^60^, its role in PDAC CAFs is still unclear.

Finally, since intra– and inter-tumour heterogeneity is closely coupled with treatment resistance and therapy response across cancers^6,30,61,62^, we investigated whether heterogeneity in the granular cell states depicted by our gene programs was conserved between our discovery and validation datasets. We calculated gene program inter-patient heterogeneity and intra-patient heterogeneity **(Methods)** and found inter– and intra-patient program loading heterogeneity varied greatly across cell types and within cell types (**Supplementary Fig. 6A, B**). Gene programs of CD4 T cells, CD8 T/NK cells, epithelial cells, and macrophages showed wide ranges of intra-patient heterogeneity across patient cohorts (**Supplementary Fig. 6B**). Across cell types, we found inter-patient heterogeneity tended to agree between the discovery and the validation datasets, except for T cell programs and macrophage programs (**Supplementary Fig. 6B**). In short, we showed in single-cell data for the first time that both intra– and inter-patient cell state heterogeneity is conserved between discovery datasets and validation datasets (intra-patient heterogeneity: *ρ* = 0.67, *p* = 1&10^-08^, inter-patient heterogeneity: *ρ* = 0.44, *p* = 0.00053). This suggests that the abundance of granular gene programs within the TME can serve as a robust estimate of heterogeneity and help guide target discovery efforts that specifically attempt to cover the entire phenotypic space.

### Co-occurring cell type-specific gene programs are highly reproducible in PDAC

We next evaluated the extent to which gene program co-occurrence within individual tumours could be reproducibly quantified to understand if fine-grained cellular subtype co-occurrence can be robustly estimated from scRNA-seq data. We computed co-occurrence of all possible gene program pairs (N=1770 total, **Methods**) separately in the discovery and the validation groups (**Fig. 2A**). To estimate this co-occurrence, we computed the Spearman correlation of the activation of each pair of gene programs across samples. This lets us estimate both the co-occurrence of gene programs within major cell types (such as if exhausted and proliferative CD8 T cells co-occur) as well as the co-occurrence of gene programs between major cell types (such as if exhausted CD8 T cells and inflammatory cancer associated fibroblasts co-occur). We quantified reproducibility of these estimates by correlating the co-occurrence in discovery and validation settings, finding highly concordant estimates (ρ=0.46, p<2×10^-16^) (**Fig. 2B**). We next used the aggregated program pair co-occurrence data to derive the positive predictive value (PPV) treating the validation co-occurrence as the ground truth and discovery as the prediction. We found that overall, the program co-occurrence PPV 80.6% for intra-cell-type co-occurrence and 74.7% for inter-cell-type co-occurrence (**Fig. 2B**). In addition, most programs saw a PPV above 50% when examined individually (**Fig. 2E**), allowing for high confidence interpretation of co-occurrence relationships.

**Figure 2:**
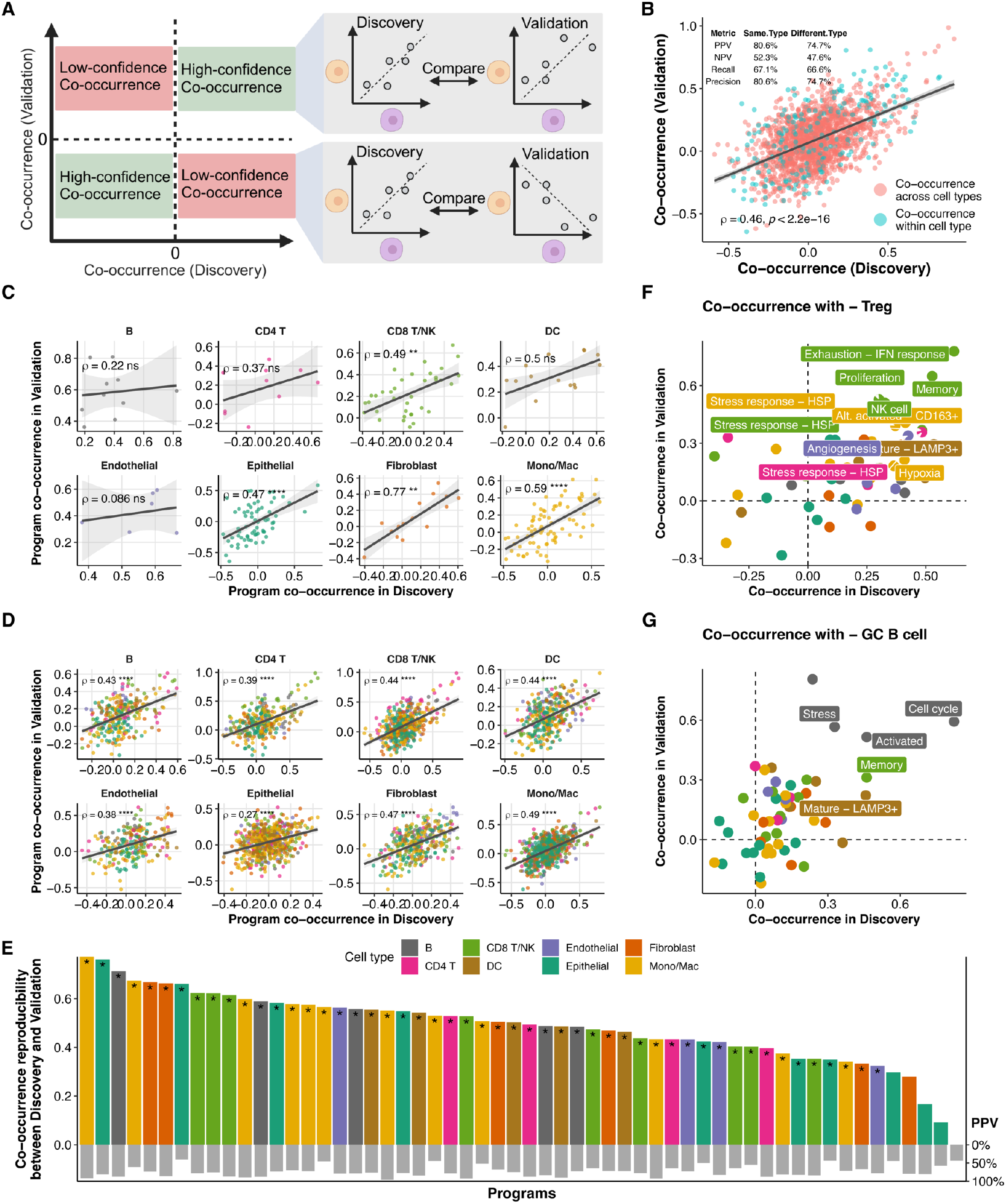
Pairwise co-occurrences of gene programs are highly reproducible. **A**. Two examples of gene program co-occurrence patterns. The program co-occurrence scores for each program pair are calculated in the discovery and the validation group separately for comparison between datasets. **B**. Program co-occurrence agreement between discovery and validation. Each dot denotes one gene program pair. Co-occurrence per program pair was measured as Spearman correlations between sample level loading means. **C**. Within cell type gene program co-occurrence agreement between discovery and validation. **D**. Across cell type gene program co-occurrence agreement between discovery and validation. **E**. Gene programs ranked by co-occurrence agreement between discovery and validation. Programs whose co-occurrence patterns are more similar between discovery and validation have higher co-occurrence reproducibility. Asterisk signs denote gene programs whose co-occurrence with other programs is significantly reproducible between discovery and validation. The grey bars below indicate empirical positive predictive value for co-occurrence of the respective gene programs. **F-G**, Sample level co-occurrence between the CD4 regulatory T cell gene program and other programs (**F**), and between the germinal center B cell program and other programs (**G**).

Next, we examined the reproducibility of cell state co-occurrence within and between major cell types. For co-occurrence of programs from the same cell type, we found highest reproducibility in fibroblasts (ρ=0.77) followed by monocyte/macrophage (ρ=0.59) (**Fig. 2C**). Gene program co-occurrence within T cells, DCs, and epithelial cells exhibited lower reproducibility between discovery and validation (ρ=0.37 to 0.5), while endothelial cells and B cells saw the lowest reproducibility across datasets (ρ=0.086 and 0.22 respectively). Nevertheless, B cell programs had positive co-occurrence in discovery (ρ=0.2-0.81) and validation (ρ=0.38-0.8), demonstrating that the overall effect size was consistent (**Fig. 2C**). We found all cell types exhibit significantly reproducible inter-cell-type program co-occurrence, most notably once again in Fibroblasts (ρ=0.47) and Monocyte/Macrophage (ρ=0.49) (**Fig. 2D**). Indeed, on a program-by-program basis we found most to have significantly reproducible inter-cell type co-occurrence relationships (**Fig. 2E**).

We next focused on the co-occurrence of specific cellular subtypes that are either significantly reproducible between discovery and validation or *a priori* known to hold clinical significance. The co-occurrence of regulatory CD4 T cells and exhausted CD8 T cells in the PDAC microenvironment was well known from mouse studies^63^, but is yet to be demonstrated in humans. We found that the CD4 T_reg_ program consistently co-occurred with the CD8 T cell exhaustion program in patient tumours (**Fig. 2F**). Also, the CD4 T_reg_ program co-occurred with the *CD163*+ alternatively activated macrophage program and the endothelial angiogenesis state program marked by *TIE1, INSR*, and *NOTCH4*. This angiogenesis program has been found to be an ECM remodeling and angiogenic signature in lung and breast cancers^64,65^ and likely interact with TAMs and regulatory T cells in breast cancer to promote an immunosuppressive phenotype in the microenvironment^65^. This T_reg_-T_exh_-Mac_alt_ co-occurrence pattern is spatially confirmed in multiple cancers^66^, suggesting our subtype program co-occurrence may be associated with spatial ecosystems in the PDAC.

Finally, we investigated whether known multi-cellular structures in the TME were consistent with our co-occurrence results. Specifically, we focused on Tertiary lymphoid structures (TLSs), localized immune aggregates within the TME that can lead to production of tumour-specific antibodies. TLSs in PDAC are usually composed of proliferating, activated, germinal center (GC) B cells, antigen presenting cells, and T cells and their presence is predictive of patient survival^67^. Therefore, we examined the co-occurrence pattern for our germinal center B cell program. Indeed, the B cell activated and cycling programs are among the top programs that co-occur with the GC program (**Fig. 2G**). In addition, we found significant co-occurrence of the activation and memory program of CD8 T cell programs with the GC B cells (ρ=0.3 in the validation group) along with mature and plasmacytoid dendritic cells (ρ=0.45). Examination of the patient sample level loadings of the TLS related cell state programs suggests variable presence of TLS in PDAC tumours (**Supplementary Fig. 7A, B**). These results suggest that in our datasets, TLS presence and phenotype could be read out from scRNA-seq. Our results imply co-occurrence of granular cellular subtypes from scRNA-seq can be meaningfully quantified from large scale atlases, and complex immune cell clusters can be identified and characterized without the need for additional spatial transcriptomics or proteomics.

### A probabilistic model to infer recurrent multi-cellular ecosystems in the TME

Beyond the pairwise co-occurrence of gene programs, multi-cellular ecosystems play a fundamental role in the TME, as these systems often act in concert, and their function is highly synergistic^68^. For this reason, we asked whether it was possible to infer ecosystems reproducibly from scRNA-seq. However, there are no existing computational tools to do this from scRNA-seq using a set of biologically meaningful gene programs as input. Therefore, we developed a novel probabilistic model (**Methods**) that can take gene programs representing cellular subtypes and infer multiple multi-cellular ecosystems, reporting each both in terms of the subtypes that co-occur as well as the ecosystem abundance within each patient sample. Importantly, our model carefully decouples the fact that programs can co-vary within the same cell (intrinsic variation) but also between cells (extrinsic variation) (**Methods**). Using maximum a posteriori inference (**Methods**), we fitted the ecosystem model separately on the discovery and validation groups using all the 50 cell type specific gene programs. We quantified four ecosystems, a number which is guided by sub-TMEs found with histology^46^.

We first contrasted the ecosystems found in the discovery and validation datasets to both validate the overall methodology and generalizability of the findings. We characterized the ecosystems by examination of the abundance of gene programs within each (**Fig. 3A**). We found significant positive correlation in 3/4 ecosystems between discovery and validation (**Fig. 3B**). We identified a Classical ecosystem enriched for corresponding epithelial programs, endothelial cells, iCAFs, and tumour associated macrophages. A Basal-like ecosystem was enriched for epithelial EMT programs, myCAF programs, and alternatively activated macrophage programs. We termed the third ecosystem immune activated as it was enriched with functional immune cell programs related to anti-tumour response, such as the memory CD4 T cell program and the effector CD8 T cell program. Notably, the TLS related programs were also enriched in the immune activated ecosystem (**Fig. 3A**). For confirmation, we found the classical and basal-like ecosystems in individual tumour samples were significantly anti-correlated in both discovery and validation cohorts (ρ = –0.43 and ρ = –0.51 respectively) (**Supplementary Fig. 8A**).

**Figure 3:**
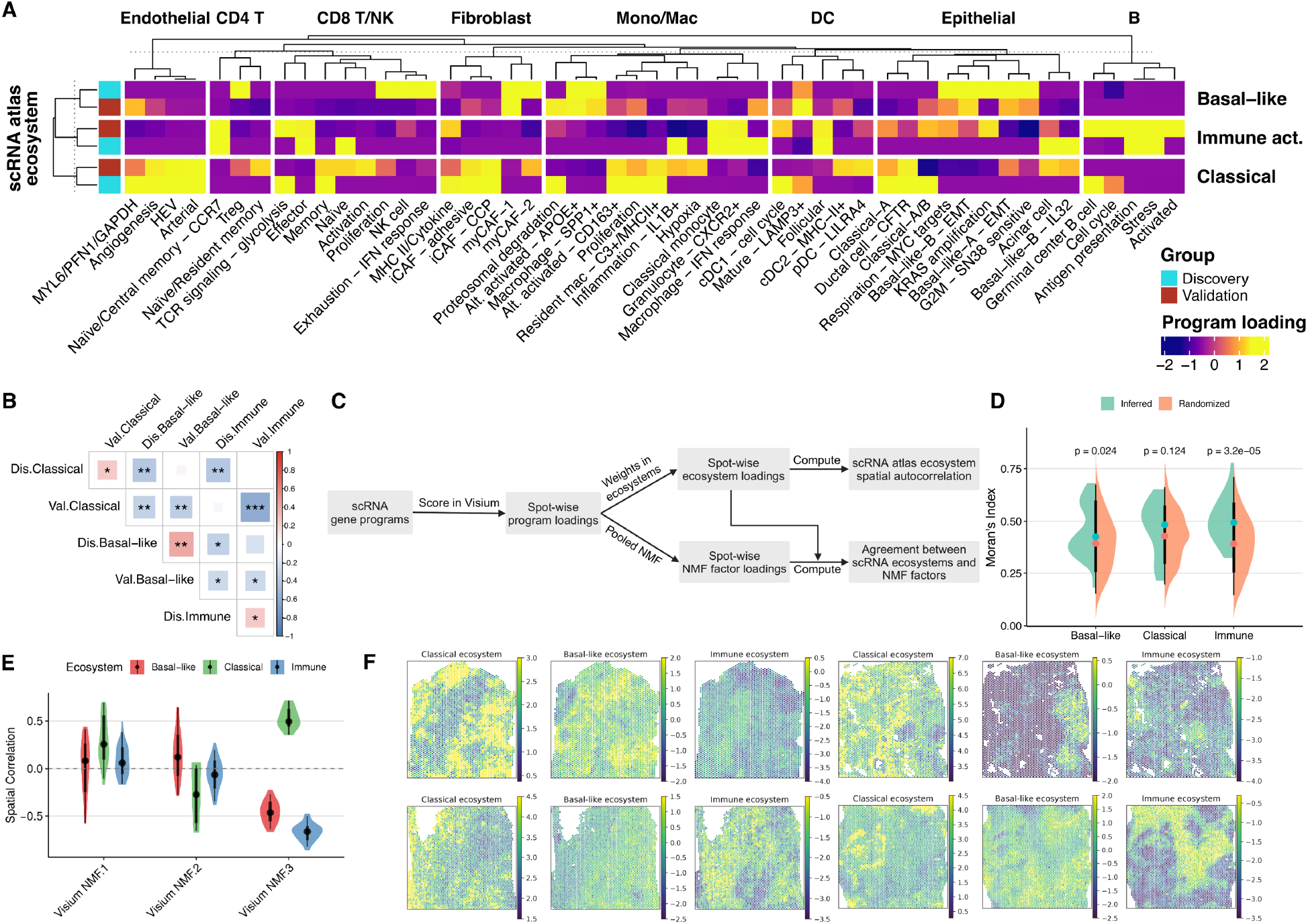
Multicellular ecosystems discovered by a probabilistic model reflect spatial variation. **A**. Gene program presence in validated ecosystems. **B**. Correlations of ecosystems between discovery and validation datasets. **C**. Workflow to validate scRNA atlas ecosystems using Visium data. **D**. Spatial autocorrelation of validated scRNA-seq inferred ecosystems compared to randomly generated ecosystems. **E**. Spot-wise correlation between scRNA ecosystems and NMF factors computed using spot-wise gene program loadings. **F**. Examples of ecosystem loadings in tumor samples.

We next investigated whether our tissue ecosystems represent spatial variation within the tumour rather than simply per-patient co-occurrence. Leveraging 24 PDAC 10x Visium spatial transcriptomics slides across 16 patients^4^, we scored individual spots for each ecosystem (**Fig. 3C, F, Methods**). We computed Moran’s I—a measure of spatial autocorrelation—for each ecosystem, then shuffled gene program weights to create 100 randomized ecosystems. We found the observed spatial autocorrelation from our scRNA-seq inferred ecosystems was higher on average than random across all three ecosystems, and significantly so for two (Basal-like: p = 0.024, immune activated: p < 0.0001, **Fig. 3D**). We also compared our ecosystems to an unsupervised learning approach for TME community discovery by generating consensus multi-program factors from spot-wise program scores using NMF. We compared the spot-wise spatial loadings of the factors with our ecosystem by computing a correlation coefficient. We found the NMF factors were not able to reproduce the Basal and immune activated ecosystem, while only the NMF factor 3 was showing moderate agreement with the classical ecosystem (**Fig. 3E**). Indeed, examining the spatial presence of the ecosystems inferred from scRNA-seq across four Visium slides shows pronounced spatial heterogeneity (**Fig. 3F**).

### Associating ecotypes with tumour genetics and clinical outcomes

To quantify the association between both our gene programs and ecosystems with tumour genetics and clinical outcomes, we curated clinical and genome sequencing data for two cohorts of PDAC tumour samples with paired bulk RNA-seq data, which we refer to as the Chan cohort^67^ (N=183 patients) and TCGA pancreatic cancer cohort^69^ (N=151 patients). We scored each sample for each of the gene programs we identified (**Methods**)^70^ and subsequently associated each sample with each ecotype by quantifying weighted combinations of gene programs (**Methods**). We first examined associations between the novel gene programs we identified and overall patient survival. Interestingly, the KRAS amplification epithelial program was significantly associated with poor patient survival in both the TCGA cohort and the Chan cohort (**Fig. 4A**). Importantly, the presence of this program was more predictive of patient survival than well the established classical and basal gene expression subtypes (**Fig. 4B**). Together, this implies such workflows that exploit high-confidence gene programs identified from single-cell RNA-seq data and validated via discovery-validation splits may be more clinically relevant than existing subtypes identified from bulk profiling approaches.

**Figure 4:**
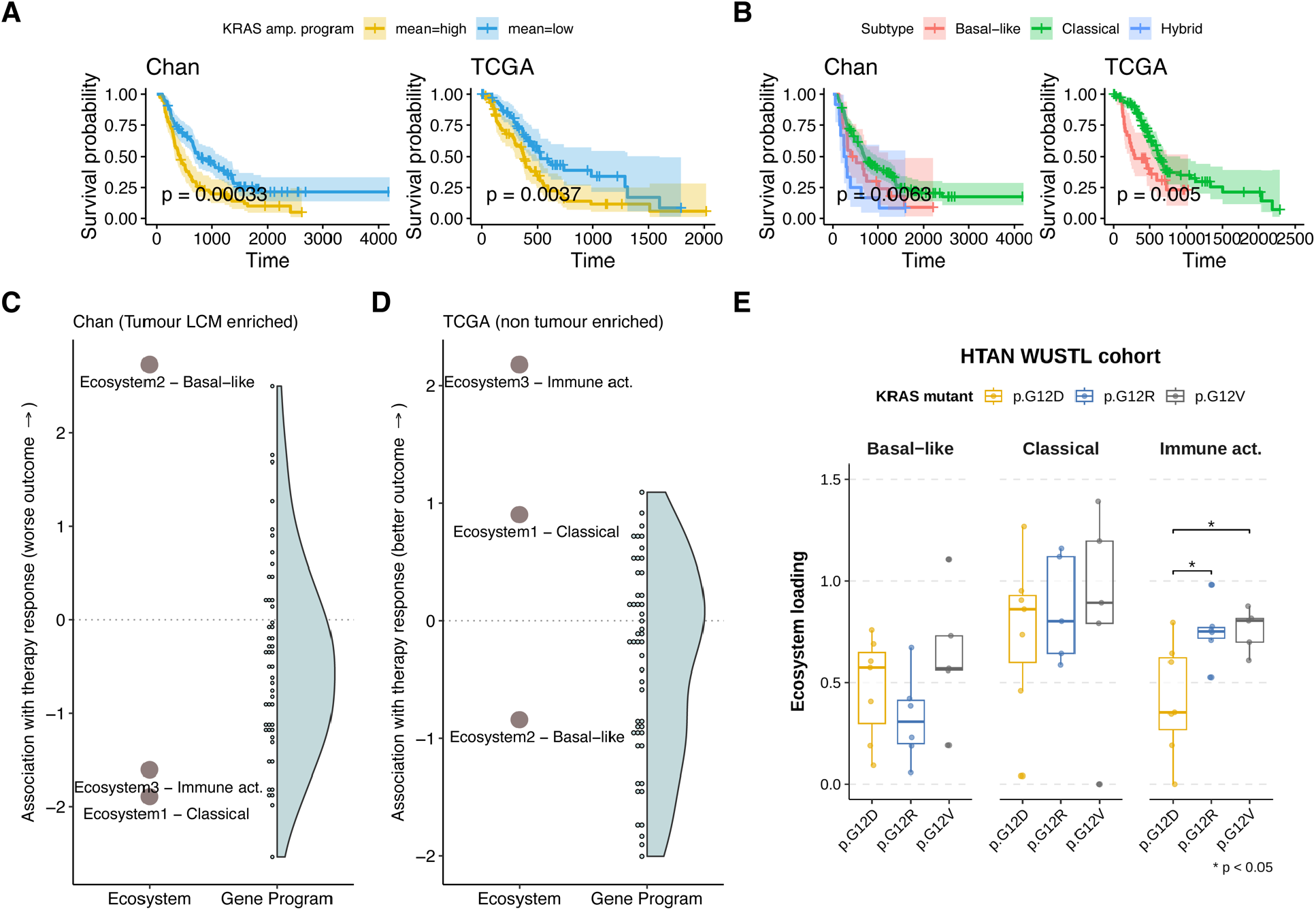
Multicellular ecosystems and cell type-specific programs are associated with patient outcomes and tumour genetics. **A**. The KRAS amplification epithelial program predicts patient survival in the Chan cohort (left) and the TCGA cohort (right). **B**. Dichotomization of patient survival by expression subtypes in the Chan and TCGA PDAC cohort. **C**. Association between ecosystems/programs and therapy outcome in the Chan cohort. **D**. Association between ecosystems/programs and therapy outcome in the TCGA cohort. **E**. Relationships between KRAS mutant and ecosystem loading in tumour samples. ‘*’: p-value < 0.05; ‘**’: p-value < 0.01, ‘***’: p-value < 0.001 (all Benjamini-Hochberg FDR corrected).

Next, we examined links between the multicellular ecosystems and patient response to therapy. Interestingly, the Basal-like ecosystem was positively associated with progressive disease after chemotherapy in both the Chan cohort (p<0.01) and the TCGA cohort (p=0.40) (**Supplementary Fig. 8B, 9**), and more significantly associated with progressive disease than any individual gene program including all Basal epithelial programs in the Chan cohort (**Fig. 4C, D**). Similarly, the immune activation ecosystem was positively associated with partial remission and negatively with progressive disease in the TCGA cohort (p<0.05) more significantly than any individual gene program (**Fig. 4C, D**). Similarly, this ecosystem was negatively associated with progressive disease in the Chan cohort, albeit not statistically significantly, (p=0.11) (**Supplementary Fig. 8B, 9**). These differences in statistical power between cohorts likely reflect differences in sample preparation: the Chan cohort underwent laser capture microdissection for the tumour epithelium, making it better placed to capture tumour associated ecosystems (Basal-like), while the TCGA cohort was not tumour enriched, which given the low cellularity of PDAC tumours (average = 35%) makes it more adept at capturing immune populations.

Finally, we contextualized the link between multicellular ecosystems and tumour genetics. PDAC tumours carrying *KRAS*-G12D mutations are associated with poorer prognosis^71^, and strategies targeting specific *KRAS* neoantigens are being developed^72^. However, the mechanisms through which different mutations affect the microenvironment in human PDAC remains unclear. We obtained *KRAS* variant calls for the HTAN Zhou (WUSTL) patient dataset^4^ in our validation cohort and fitted a linear mixed model between ecosystem presence and *KRAS* variant status (**Methods**). We found that compared to *KRAS*-G12D, tumours with a *KRAS*-G12V/G12R mutations had a significantly higher presence of the immune activated ecosystem, while tumours with *KRAS*-G12R mutations had a reduced presence of the basal-like ecosystem (**Fig. 4E**). Importantly, we retrieved similar *KRAS* variant calls in the bulk Chan patient cohort^69,70^ and found consistent effect sizes for all associations (**Supplementary Fig. 8C, Supplementary Table 3**), validating the overall findings. The decreased presence of the immune activated ecosystem in *KRAS*-G12D PDAC suggests differences in anti-tumour immune response may underline prognosis and effectiveness of immunotherapy^71,72^. Together, these results suggest that the specific driver mutations in PDAC have a significant effect on the phenotypes and cell communities in the TME.

## Discussion

Transcriptome profiling of tumour samples has historically been used to identify molecular markers that could facilitate cancer diagnosis, prognosis prediction and identification of molecular candidates to target therapeutically^73–76^. The advent of single cell technologies has facilitated the identification of the relative distribution of cell populations specific for certain cancer types, predicting clinical outcomes and uncovering specific cellular subtypes as targets^77–80^. However, it is becoming increasingly clear that focusing on the expression pattern of a handful of genes or the presence/absence of certain cell populations is not sufficiently informative to gain useful biological insights into the whole TME, even though this information could be leveraged to design targeted therapies aimed at tackling the combined action of multiple cell types. In this context, spatial information is extremely useful to identify co-localization of different cell types, whose crosstalk requires a certain three-dimensional organization to synergistically alter tumour growth and survival. However, a substantial number of samples are needed to robustly identify co-occurrence of interacting cells across patients, especially considering that the cellular subtypes of interest are potentially rare. Therefore, the current cost and/or limitations of spatial transcriptomics and proteomics limit the identification of reproducible co-existing cell states with prognostic value^79^. By contrast, the number of scRNA-seq datasets from TMEs that contain transcriptome-wide expression quantification at single-cell resolution is increasing exponentially, allowing, in theory, to identify co-occurrence of granular cell states and associate them with clinical outcomes.

Here we have undertaken a unique discovery-validation analysis on the largest PDAC scRNA-seq atlas to-date to quantify the extent to which granular cellular subtypes and ecosystems can be reliably inferred. Our 50 reproducible cell type specific gene programs captured previously known heterogeneity such as classical and basal tumour transcriptional but also novel subtypes such as two iCAF subtypes as well as a novel *KRAS amplification* epithelial cell program significantly associated with survival. We further revealed for the first time that co-occurrence of gene programs can be reproducibly quantified from large scRNA-seq datasets. We further identified a TLS pattern centered around the GC B cell program, recapitulating a spatial structure important for anti-tumour immune response in multiple cancers including PDAC^81^. We introduced a new model for inferring ecosystems in TME and found reproducible ecosystems, which were validated in spatial transcriptomic data and associated with both driver gene mutants as well as response to therapy.

Overall, we show that large scRNA-seq atlas can be employed to not only identify TME cellular subtypes and ecosystems with high confidence, but that these have profound biological and clinical significance, as the frequency of co-occurrence of certain programs and the presence of certain ecosystems are associated with patient survival. These analyses suggest that in the future, scRNA-seq datasets will be used to study the broad interplay among pro-tumorigenic cell populations in the TME. This approach may eventually expand the clinical usability of single cell data, offering a new lens to develop innovative therapeutic strategies targeting PDAC.

Despite these findings, our study had multiple limitations. While our discovery group was well balanced between four datasets, the validation cohort was heavily weighted towards one cohort, which may have limited the cell state program validation process though we note that, since there was no overlap in samples, it still functions as a well-powered validation cohort. In addition, the lack of well-standardized and curated clinical metadata across the single-cell cohorts limited the ability to directly associate cell states and ecosystems with clinical covariates, instead necessitating mapping them to bulk datasets with standardized metadata.

Moving forward, we identified several venues to extend the project. Future spatial profiling of the PDAC microenvironment at cellular resolution in large cohorts will help to create a more comprehensive understanding of the cell state co-occurrence related to therapy resistance and disease progression. Integrating T cell receptor (TCR) or B cell receptor (BCR) sequencing analysis will help to delineate relationships between lymphocyte clonality with functional cell states, potentially revealing specific anti-tumour T cell clones and effects of therapy on the tumour immune cell population. In the long term, our analysis workflow may be adapted to multiple cancer types, leveraging the ever-increasing tumour single-cell data space to create a pan-cancer microenvironment cell state atlas, and identifying clinically relevant cellular subtypes across cancers. The increasing characterization of the TME at single-cell level will enable the identification of concomitant presence of multicellular ecosystems in various types of cancer. Our approach unlocks a new way to analyse scRNA-seq data that will not only facilitate prognosis and assessment of therapeutic response, but will provide valuable information to design specific therapies aimed at rewiring the milieu compatible with the co-existence of pro-tumorigenic cellular subtypes and ecosystems.

## Methods

### Data curation

A total of 8 publicly available PDAC single-cell RNA-seq (scRNA-seq) datasets, which include 214 samples, were curated^4,32–37,47^. Most datasets were available for downloading through the Gene Expression Omnibus database. One dataset was available on author consent, in which case we contacted the authors for permission. Each dataset contains varying types of samples. We excluded samples which are of the following nature: metastasis, normal pancreas, peripheral blood, and adjacent normal tissue. In total, 163 samples labeled as primary PDAC tumour were retained for downstream analysis.

After sample filtering, we performed in house pre-processing and quality control given all datasets were available in the CellRanger output format. Doublet scores for single cells were calculated and their doublet status was called using doubletFinder_v3 from DoubletFinder v.2.0.3^82^. Cells labeled as doublets were removed from each sample. After doublet removal, cells with mitochondrial RNA content percentage greater than 20%, detected number of genes lower than 500, and total transcript counts lower than 1000 were removed as low quality cells. Finally, the UMI counts were normalized by per-cell size factor and log2-transformed with logNormCounts from scuttle v.1.8.4^83^.

### Cell annotation unification

We performed re-annotation to unify the cell annotation in each patient cohort in our curated dataset. First, we obtained a single-cell PDAC atlas built from divergent and heterogeneous datasets^81^. The ‘celltype2’ annotation from the atlas was selected as the reference labels after comparison of resolution and quality of all annotation levels in the atlas. Then, we performed unbiased cell type recognition at the single cell level using SingleR from SingleR v.2.0.0^84^. Briefly, SingleR first trains a classifier on the PDAC atlas reference dataset with reference labels by leveraging differential expression between labels, then the labels were assigned to each single cell in the query dataset using the classifier combined with iterative fine-tuning. We ran SingleR for each patient cohort in the curated dataset independently with the following settings: excluding ribosomal and mitochondrial genes in the reference, selecting top 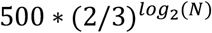 (N = # unique genes) DE genes with ranking by log-fold change, aggregating the reference into pseudo-bulk samples with default settings, and enabling fine-tuning with default settings. After annotation by SingleR, quality checks were performed by examining the SingleR predicted scores for each label in single cells in each cohort, the distribution of delta between assigned label score and median score across labels, and the distribution of scores for each label (**Supplementary Fig. 1C**). Final validation was done by examining known cell type marker expressions in each assigned cell type. Finally, the SingleR pruned.label column was selected and cell type labels were assigned as the unified annotation for single cells. To facilitate downstream analysis, endocrine cells and neural cells were removed from the curated dataset, and acinar and ductal cells were both labeled as epithelial cells. The eight consolidated cell type labels were then shortened for readability.

### Discovery-Validation data split

We first subset the full curated dataset by cell annotation, splitting it into eight cell type subsets: Epithelial cell, Endothelial cell, Fibroblast, Monocyte/Macrophage, mDC, CD4 T cell, CD8 T/NK cell, and B cell. We then split each cell type subset into two 4-cohort groups, one as the discovery group and the other as the validation group. The discovery group contained 74 samples from the Peng, Steele, Werba, and Chan patient cohorts, while the validation group contains 89 samples from the Chen, Lee, Lin, and Zhou patient cohorts. In total, the discovery-validation split resulted in eight discovery-validation pairs of cell type subsets, with each pair containing single cells assigned to one cell type.

### Reproducible analysis pipeline

We built a comprehensive analysis pipeline to generate reproducible results from our curated dataset and reduce human-introduced variations in analyses across the cell types and between the discovery group and the validation group. We used Snakemake v.7.25.0^85^ for parameter and input-output control for all analysis scripts and visualization steps.

### Defining a set of robust lineage-specific cell state gene programs

We carried out integrative NMF (iNMF) using LIGER^82^ for the discovery and validation group of patient cohorts across cell types independently to generate cell lineage-specific gene programs that capture the phenotypic heterogeneity within each cell type in the PDAC microenvironment. LIGER runs iNMF to decompose the expression matrices from multiple batches (patient cohorts) and remove batch effect at the same time by maximizing a basis matrix shared between cohorts while minimizing the multiple cohort-specific basis matrices that represent patient cohort level biases. We defined the batches to be patient cohorts to focus on capturing heterogeneity present across tumours through cell lineage-specific programs. We followed the suggested workflow of LIGER to perform iNMF. Importantly, the following steps were performed after the discovery-validation split as well as an 8-way cell type split within the discovery and the validation group, leading to 16 independent gene program extraction steps, 8 independent program validation steps, and 8 independent program merging steps.

### Program extraction

First, we selected variable genes from non-ribosomal/mitochondrial protein-coding genes using from rliger v.1.0.0^83^ with default settings. We then created a liger object using the raw counts matrix, removed genes with zero expression, and normalized and scaled gene expression following suggested workflow from rliger documentation before performing a subset to the variable genes. Finally, after filtering to variable genes, we fitted iNMF on the expression matrix using optimizeALS. We set rank (k) to 20, 10, 5, 10, 10, 20, 10, and 10 for program extraction from epithelial cell, fibroblast, endothelial cell, CD4 T cell, CD8 T/NK cell, monocyte/macrophage, mDC, and B cell respectively. We set lambda to 5 for all cell types as we did not find it to be a major factor affecting extraction results. The gene loadings in each program were obtained as the W matrix from the fitted liger object, and the program loadings in single-cells were obtained as the H.norm matrix from the fitted liger object.

### Program validation

Our program validation process was performed between discovery and validation for each cell type. To start, we transformed the discovery and validation gene loading (W) matrix with log1p in R. We then computed the spearman correlation and Minkowski distance (with p = 1) between all possible pairs of discovery and validation programs within the cell type. The Hungarian algorithm was used to generate a one-to-one relationship between discovery and validation programs by permuting the rows of the distance and correlation matrices to minimize and maximize the trace of the matrices respectively. After assigning the one-to-one partner of each discovery program, we rejected any discovery programs whose gene loading correlation with their validation program partner was below 0.5, or if the gene loading distance with their validation program partner was above 25,000. We further confirm validated programs by permutation test. Gene loadings in each validated program were permuted to generate a 1000-sample test set and the correlation and distance to the validation program partner were calculated for fitting a null distribution. A discovery program is finalized as a validated program if the real distance and correlation are laid beyond 1% of the null distribution.

### Program merge

We collapsed validated programs which were too similar to each other given NMF does not guarantee orthogonality between the extracted factors. We looked at all possible pairs of validated programs within each cell type and marked any pair whose gene loading correlation was above 0.4 for merging. The merging step leveraged simple linear models. For a pair of validated programs, e.g. sig-1 and sig-2, marked for merging, we first computed their explained expression y by multiplying program loading in cell and gene loading in program respectively. Then, we obtained the merged program’s gene loading x by taking the mean of gene loadings in sig-1 and sig-2. Finally, the single-cell loading of the merged program was obtained by fitting lm(y ∼ 0 + x) and pulling the coefficients. After merging, the newly merged program was included for downstream analysis while sig-1 and sig-2 were excluded.

### Program rescoring in the validation group

Using the robust set of validated and merged cell type specific programs, we rescored the validation group single cells to get single-cell program loadings that can be compared with program loadings in the discovery group in downstream analysis. The rescoring step was performed for each cell type and each patient cohort independently, leading to 32 rescoring steps.

To start, gene expression in a validation group patient cohort was normalized using runSeuratNormalizeData from singleCellTK v.2.8.0^86^ with default settings. Then, for rescoring for programs of, for example, cell type A, we subset an expression matrix to cells annotated as cell type A and variable genes used in the program extraction step. Finally, we performed non-negative least squares to fit a simple linear model to the subset validation expression with cell type A program gene loadings, and pulled out the estimated optimal coefficient matrix as the rescored program loading in single-cells in validation. We used fcnnls from NMF v.0.26^87^ with default settings for this step.

### Program contextualization with known cell states

We combined multiple approaches to get a comprehensive understanding of the biological meaning of our validated cell type-specific programs. We curated multiple PDAC focused malignant, stromal, and immune cell state marker lists from literature (**Supplementary Table 4**). To associate the programs with the known cell states, we selected the top 50 loaded genes of each program and performed a hypergeometric test with the cell state marker lists from its respective cell type, using GeneOverlap v.1.34.0^88^ in R. The human reference hg19 genome^89^ was used as the background for significance testing. We made a preliminary biological interpretation of the programs based on significance (P-value < 1e-3) and high Jaccard index with cell state marker lists. Then, the programs were further subjected to GO and KEGG overrepresentation analysis using the clusterProfiler v.4.6.2 package^90^. Specifically, the biological process subcategory of GO and KEGG PATHWAY gene sets were chosen for the analysis. The top ten enriched gene sets in each category were used to help correct or confirm our initial program interpretation. After the first correction, we also applied GSEA to the programs with gene sets mined from the Curated Cancer Cell Atlas (3CA)^91^ to resolve any remaining uncertainty. We ran fgsea from fgsea v.1.24.0^92^ with default settings, while setting nPermSimple to 10,000, as suggested by the documentation. We used the top 10 positively enriched 3CA gene programs to guide a second program interpretation correction step. Programs that were not associated with known cell states previously were further characterized after this analysis. Finally, we completed a manual check across programs by looking at the top 10 loaded genes, focusing on the programs that were yet to be contextualized and searching in the literature for novel evidence of their biological meaning in PDAC. Any novel programs remained were named by their top loaded genes.

### Ambient RNA program removal

Due to technical limitation of droplet-based single-cell RNA-seq experiments, some reads associated with a cell barcode may originate from ambient RNA^54^, contributing to background noises that are picked up by the granular cell state gene programs. Therefore, we denoted gene programs in a cell lineage that are marked by lineage markers from other lineages to be ambient RNA programs and removed such ambient programs from downstream analysis.

### Cell state co-occurrence

To study co-occurrence between programs in the PDAC microenvironment, we constructed sample level program loading profiles by averaging program loadings across single-cells. We then calculated the correlation across samples between programs both within cell type and across cell types. By comparing program loading correlations between the discovery group and the validation group, we define program pairs whose loading correlation was above 0.4 and differed by less than 0.25 in discovery and validation as high confidence pairs that tend to co-occur with each other.

### Scoring RNA-seq samples for cell state programs and ecosystems

To score bulk PDAC tumour samples for our cell state gene programs, we performed gene set variation analysis (GSVA) on each bulk sample using gsva from GSVA^93^ ver. 1.46.0, with the top-50 loaded genes in each gene programs, and used the program enrichment scores or downstream analysis. To score bulk PDAC tumour samples for our ecosystems, we performed multiplication between the sample-by-program enrichment score matrix and the program-by-ecosystem abundance matrix, to obtain a sample-by-ecosystem enrichment score matrix.

### Association between ecosystems and KRAS variants

KRAS variant information was obtained from published data and from collaborators. To account for the patient-repeated measurements in the Chan cohort, we fitted a linear mixed model with the *lmerTest* package in R with the KRAS mutation status as the fixed effect and the patient as the random effect. For the TCGA cohort to estimate effect of KRAS variant on ecosystem enrichment scores we used a simple linear model while controlling for the expression subtype (Basal-like or Classical) of the tumour.

### Association between gene programs, ecosystems, and clinical outcomes

To associate the cell type-specific gene programs and multicellular ecosystems with clinical outcomes, we mapped scored RNA-seq samples in the Chan cohort and the TCGA cohort to metadata and clinical data. We fitted univariate linear regression models to each gene program-clinical outcome pair and each ecosystem-clinical outcome pair, and retrieved the t-statistics as estimate of association. We further fitted univariate Cox Proportional-Hazard ratio models to obtain hazard ratios for each program and ecosystem. For survival analysis, we used consolidated survival data from each cohort to plot KM curves to stratify patients.

### Cell state co-occurrence ecosystems discovery model

To further dissect the relationships between programs, we created a statistical model to deconvolve program co-occurrence and discover multi-cellular population ecosystems.

When considering why programs are (anti-) correlated in single-cells, there can be two reasons. Firstly, within single cells, programs may be correlated due to gene regulation, epigenetics, etc. We defined correlations resulting from these mechanisms cell intrinsic program correlations. Secondly, within patients, cells with a high program of one type may be present with (or exclude) cells with a different program. This could be due to paracrine signaling. We defined these correlations cell extrinsic program correlations. If, for example, there is positive correlation between loading of two programs only at the patient level, but not at the single cell level, then the correlation is dominated by cell extrinsic program correlation, and vice versa for cell intrinsic program correlation. We note that in both scenarios, the overall correlation between the programs is the same, even if the mechanisms and interpretation are different. So, it is important to distinguish the two types of correlation in our model.

### Model data definition

We start by defining *L* as the number of ecosystems we want to get, *C* as the number of cell types, *N* as the total number of cells, *K* as the total number of cell state programs, and *P* as the total number of patient samples. Therefore, we have:

The set of all cell types: ***C*** = {*C*_1_, *C*_2_, *C*_3_, … *C*_*c*}_

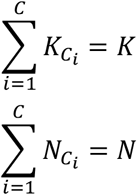

The set of all patients: ***P*** = {1, 2, 3,…, *P*}

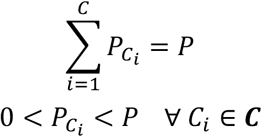

Then, for each cell type *C*_*i*_, we have 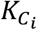 programs across 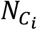 ingle-cells, and each single-cell belongs to one of 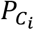 patients.

Next, we define some cell-type-specific variables. For each cell type *C*_*i*_, we have a matrix 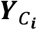 storing the cell state program loadings in single-cells:

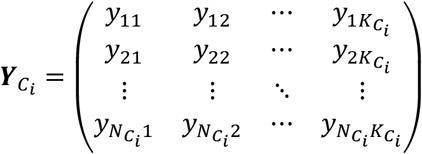

In 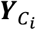 each row represent one 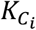 –dimensional vector for a single-cell. This vector stores the loadings of each cell state program for cell type *C*, in the cell. Of note, each single-cell is assigned to one cell type only, and thus only shows up in one of the 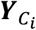 matrices.

To further indicate the patient sample each single-cell belongs to, we introduce a sample indicator vector for each cell type *C*_*i*_:

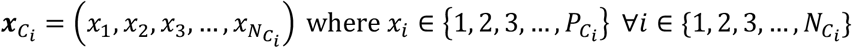

### Model parameter definition

We first introduce the ecosystem parameters by defining:

1. The ecosystem loading matrix ***H*** = (*h*_*ij*_)_*P*×*L*_
2. The ecosystem basis matrix ***W*** = (*w*_*ij*_)_*L*×*K*_
3. The patient level program loading estimate matrix considering extrinsic covariance ***Φ*** = (***ϕ***_*ij*_)_*P* ×*K*._ = ***H*** × ***W***

Where each row *w*_*l*_ of ***W*** is a *K*-dimensional vector of program weights in ecosystem *l*, and each row *h*_*p*_ of ***H*** is an *L*-dimensional vector of ecosystem weights in sample *p*.

With the ecosystem related parameters defined, we introduce:

1. The patient level program loading estimate matrix considering intrinsic + extrinsic covariance ***M*** = (*μ*_*ij*_)_*p*×*K*_
2. The cell-type-specific patient level program loading sub-matrices 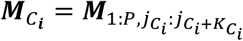 Where 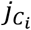 is the index for the first program of cell type *C*_*i*_ in ***M***, specifically:

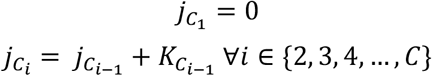

We lastly explicitly define, for each cell type *C*, the cell intrinsic program covariance matrix as:

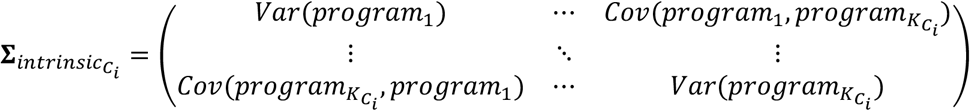

### Model definition

First, we let

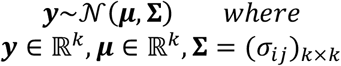

We then consider cell extrinsic and intrinsic program correlations separately.

Cell extrinsic program correlations are correlations in the means of the program loadings for each patient. We thus introduce *P* mean vectors ***μ***_*p*_ (one ***μ***_*p*_ per patient) that are each *K*-dimensional and encode the mean program loading of patient *p*. Of note, Then, we can write:

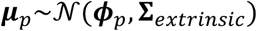

Here, ***ϕ***_*p*_ is the prior on the patient mean program loading, and **Σ**_*extrinsic*_ is the program by program *K* × *K* covariance matrix of cell extrinsic program correlations.

Cell intrinsic program correlations are the correlation of the programs conditioned on the mean program loadings of each patient. We thus introduce *N* cell level program loading vectors 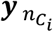 that are each 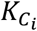 –dimensional and encode the program loadings for cell 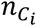 where *C*_*i*_ ∈ **C**.

Therefore, we can write:

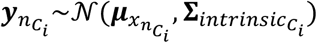

Here, 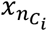 selects the patient sample corresponding to cell *n*, so 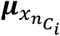 corresponds to the mean program loadings of the patient corresponding to cell 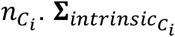 is the *K* × *K* covariance matrix that denotes the program correlation after conditioning on the patient means. Using Gaussians allows us to interpret the total covariance as the sum of the intrinsic and extrinsic covariance, i.e. 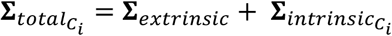.

Together, we can write the hierarchical model:

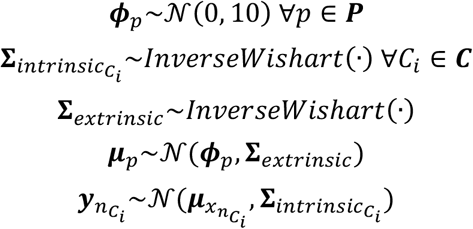

Here,

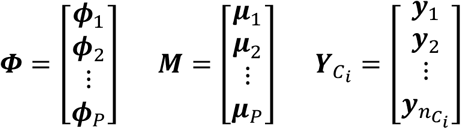

### Target function

The target function of the model:

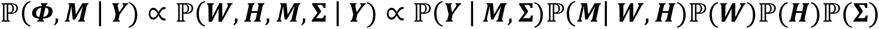

The joint posterior is then:

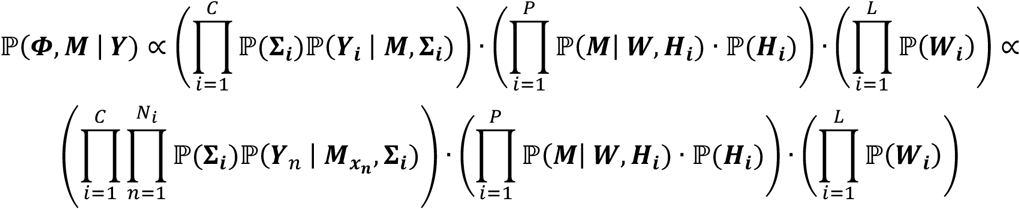

To encourage the model to find more distinct ecosystems, we add an orthogonality term to the model target function. The upper triangular matrix of the transpose cross product of the ecosystem basis matrix ***W*** is used to calculate a mean as a proxy for ecosystem similarity. This term is weighted by a hyperparameter *λ* and minimized in the target function. Therefore, using maximum a posteriori inference, we maximize the new target function including the log joint posterior and the orthogonality term:

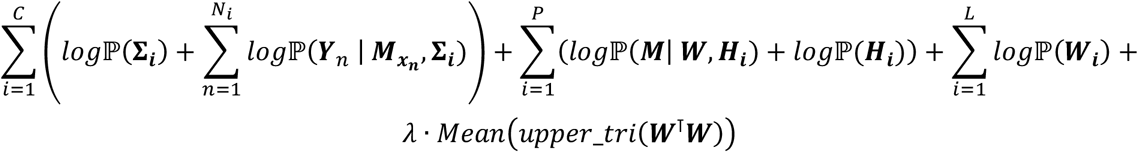

### Data preparation for the ecosystem discovery model

To increase the power of our model, we removed patient samples which contained fewer than 20 single-cells. To speed up the inference process, we sampled down each patient sample to 1500 single-cells. For each cell state programs, we scaled the program loadings in single-cells across all cells to get z-scores as input to the model.

### Model initialization and optimization

We coded the ecosystem discovery model using the statistical programing language Stan. We used cmdstan_model from the R package cmdStanR ver. 0.5.3^94^ to compile the model. Then, we used optimize from cmdStanR to initialize and fit the model to our data, with settings: algorithm=“lbfgs”, tol_param=1e^-7^, and iter=8000. The model parameters were initialized as specified below.

Model initialization was provided for **Σ**_*intrinsic*_ for each cell type and for ***W*** and ***H***. We performed NMF on the sample level cell state program loading mean matrix with rank=*L* to obtain the initial values of the ecosystem matrices. Covariance between program loadings within respective cell types were input as the initial values of **Σ**_*intrinsic*_

### PDAC Visium data

Visium data was downloaded from Synapse^95^ with author permission. Tumour sample slides from primary tumours were selected. Preprocessing was done using the SpatialData^96^ library. Low quality spots containing more than 20% hemoglobin genes and fewer than 1000 detected genes were filtered. Non-protein coding genes, mitochondrial genes, ribosomal genes, and hemoglobin genes were excluded from downstream analysis. Raw UMI counts were log-normalized by calling computePooledFactors followed by logNormCounts with default parameters.

### Scoring programs and ecosystems in Visium

Top 20 genes ranked by weight in each gene program were used to compute spot-wise scores in Visium PDAC samples. Since each spot on the slide is effectively a mini bulk sample, we leveraged gene set variation analysis (GSVA) to obtain an enrichment score for each program in each spot of each Visium slide. Next, program scores in each spot were scaled to 0-1 range. Consensus weighing factors for each ecosystem was obtained by removing programs whose weights had opposite signs in discovery vs. validation. We then obtained spot-wise ecosystem scores as the sum of the program scores weighted by the consensus weighing factors. Randomized ecosystem baselines were generated by shuffling the consensus weighing factors.

### Concatenated NMF

To obtained consensus NMF factors. All Visium spots were concatenated to create one spot x program score matrix. A single NMF model was fitted on the pooled matrix, and the expression was decomposed into three factors common across images.

### Spatial autocorrelation

To measure spatial autocorrelation, we computed the Moran’s Index for the inferred ecosystems, the randomized ecosystems, and the concatenated NMF factors in each Visium image. We used Squidpy^97^ graph algorithms with default parameters for Moran’s I computation.

## Data and code availability

All data and code required to reproduce this study can be found at https://zenodo.org/records/13332977 and https://github.com/camlab-bioml/PDAC_cellular_niche_analyses.

## Contributions

Project conception: CY, KC. Data curation: CY, SG, TTJ, AZ. Data analysis and coding: CY, MG, SG, GHJ, TTJ, AZ. Result interpretation and manuscript writing: CY, KC. Project direction: GO, FN, KC.

## Acknowledgements

We thank Life Science Editors for editing services (www.lifescienceeditors.com).

