## Supplementary materials for "Inferring tumour microenvironment ecosystems from scRNA-seq atlases"

Chengxin Yu *et al.*

**This PDF file includes:**

Detailed description of the cellular subtype ecosystem discovery model
Figs. S1 to S10
Legends for tables S1 to S4

**Other Supplementary Material for this manuscript includes the following:**

Tables S1 to S4

**Detailed description of the cellular subtype ecosystem discovery model**
When considering why programs are (anti-) correlated in single-cells, there can be two reasons. Firstly, within single cells, programs may be correlated due to gene regulation, epigenetics, etc. We defined correlations resulting from these mechanisms cell intrinsic program correlations. Secondly, within patients, cells with a high program of one type may be present with (or exclude) cells with a different program. This could be due to paracrine signaling. We defined these correlations cell extrinsic program correlations. If, for example, there is positive correlation between loading of two programs only at the patient level, but not at the single cell level, then the correlation is dominated by cell extrinsic program correlation, and vice versa for cell intrinsic program correlation. We note that in both scenarios, the overall correlation between the programs is the same, even if the mechanisms and interpretation are different. So, it is important to distinguish the two types of correlation in our model.

<u>Model data definition</u>
We start by defining $L$ as the number of niches we want to get, $C$ as the number of cell types, $N$ as the total number of cells, $K$ as the total number of cell state programs, and $P$ as the total number of patient samples. Therefore, we have:

$$\text{The set of all cell types: } \boldsymbol{C} = \{C_1, C_2, C_3, \ldots, C_c\}$$

$$\sum_{i=1}^{C} K_{C_i} = K$$

$$\sum_{i=1}^{C} N_{C_i} = N$$

$$\text{The set of all patients: } \boldsymbol{P} = \{1, 2, 3, \ldots, P\}$$

$$\sum_{i=1}^{C} P_{C_i} = P$$

$$0 < P_{C_i} < P \quad \forall\, C_i \in \boldsymbol{C}$$

Then, for each cell type $C_i$, we have $K_{C_i}$ programs across $N_{C_i}$ single-cells, and each single-cell belongs to one of $P_{C_i}$ patients.

Next, we define some cell-type-specific variables. For each cell type $C_i$, we have a matrix $\boldsymbol{Y}_{C_i}$ storing the cell state program loadings in single-cells:

$$\boldsymbol{Y}_{C_i} = \begin{pmatrix} y_{11} & y_{12} & \cdots & y_{1K_{C_i}} \\ y_{21} & y_{22} & \cdots & y_{2K_{C_i}} \\ \vdots & \vdots & \ddots & \vdots \\ y_{N_{C_i}1} & y_{N_{C_i}2} & \cdots & y_{N_{C_i}K_{C_i}} \end{pmatrix}$$

In $\boldsymbol{Y}_{C_i}$ each row represent one $K_{C_i}$-dimensional vector for a single-cell. This vector stores the loadings of each cell state program for cell type $C_i$ in the cell. Of note, each single-cell is assigned to one cell type only, and thus only shows up in one of the $\boldsymbol{Y}_{C_i}$ matrices.

To further indicate the patient sample each single-cell belongs to, we introduce a sample indicator vector for each cell type $C_i$:

$$\boldsymbol{x}_{C_i} = \left(x_1, x_2, x_3, \dots, x_{N_{C_i}}\right) \text{ where } x_i \in \left\{1, 2, 3, \dots, P_{C_i}\right\} \ \forall i \in \{1, 2, 3, \dots, N_{C_i}\}$$

<u>Model parameter definition</u>
We first introduce the niche parameters by defining:

1. The niche loading matrix $\boldsymbol{H} = (h_{ij})_{P \times L}$
2. The niche basis matrix $\boldsymbol{W} = (w_{ij})_{L \times K}$
3. The patient level program loading estimate matrix considering extrinsic covariance $\boldsymbol{\Phi} = (\phi_{ij})_{P \times K} = \boldsymbol{H} \times \boldsymbol{W}$

Where each row $w_l$ of $\boldsymbol{W}$ is a $K$-dimensional vector of program weights in niche $l$, and each row $h_p$ of $\boldsymbol{H}$ is an $L$-dimensional vector of niche weights in sample $p$.

With the niche related parameters defined, we introduce:

1. The patient level program loading estimate matrix considering intrinsic + extrinsic covariance $\boldsymbol{M} = (\mu_{ij})_{P \times K}$
2. The cell-type-specific patient level program loading sub-matrices $\boldsymbol{M}_{C_i} = \boldsymbol{M}_{1:P, j_{C_i}:j_{C_i}+K_{C_i}}$

Where $j_{C_i}$ is the index for the first program of cell type $C_i$ in $\boldsymbol{M}$, specifically:

$$j_{C_1} = 0$$
$$j_{C_i} = j_{C_{i-1}} + K_{C_{i-1}} \ \forall i \in \{2, 3, 4, \dots, C\}$$

We lastly explicitly define, for each cell type $C_i$, the cell intrinsic program covariance matrix as:

$$\boldsymbol{\Sigma}_{intrinsic_{C_i}} = \begin{pmatrix} Var(program_1) & \cdots & Cov(program_1, program_{K_{C_i}}) \\ \vdots & \ddots & \vdots \\ Cov(program_{K_{C_i}}, program_1) & \cdots & Var(program_{K_{C_i}}) \end{pmatrix}$$

<u>Model definition</u>
First, we let

$$\boldsymbol{y} \sim \mathcal{N}(\boldsymbol{\mu}, \boldsymbol{\Sigma}) \qquad where$$
$$\boldsymbol{y} \in \mathbb{R}^k, \boldsymbol{\mu} \in \mathbb{R}^k, \boldsymbol{\Sigma} = (\sigma_{ij})_{k \times k}$$

We then consider cell extrinsic and intrinsic program correlations separately.

Cell extrinsic program correlations are correlations in the means of the program loadings for each patient. We thus introduce $P$ mean vectors $\boldsymbol{\mu}_p$ (one $\boldsymbol{\mu}_p$ per patient) that are each $K$-dimensional and encode the mean program loading of patient $p$. Of note, Then, we can write:

$$\boldsymbol{\mu}_p \sim \mathcal{N}(\boldsymbol{\phi}_p, \boldsymbol{\Sigma}_{extrinsic})$$

Here, $\boldsymbol{\phi}_p$ is the prior on the patient mean program loading, and $\boldsymbol{\Sigma}_{extrinsic}$ is the program by program $K \times K$ covariance matrix of cell extrinsic program correlations.

Cell intrinsic program correlations are the correlation of the programs conditioned on the mean program loadings of each patient. We thus introduce $N$ cell level program loading vectors $\boldsymbol{y}_{n_{C_i}}$ that are each $K_{C_i}$-dimensional and encode the program loadings for cell $n_{C_i}$ where $C_i \in \boldsymbol{C}$. Therefore, we can write:

$$\boldsymbol{y}_{n_{C_i}} \sim \mathcal{N}(\boldsymbol{\mu}_{x_{n_{C_i}}}, \boldsymbol{\Sigma}_{intrinsic_{C_i}})$$

Here, $x_{n_{C_i}}$ selects the patient sample corresponding to cell $n$, so $\boldsymbol{\mu}_{x_{n_{C_i}}}$ corresponds to the mean program loadings of the patient corresponding to cell $n_{C_i}$. $\boldsymbol{\Sigma}_{intrinsic_{C_i}}$ is the $K \times K$ covariance matrix that denotes the program correlation after conditioning on the patient means. Using Gaussians allows us to interpret the total covariance as the sum of the intrinsic and extrinsic covariance, i.e. $\boldsymbol{\Sigma}_{total_{C_i}} = \boldsymbol{\Sigma}_{extrinsic} + \boldsymbol{\Sigma}_{intrinsic_{C_i}}$.

Together, we can write the hierarchical model:

$$\boldsymbol{\phi}_p \sim \mathcal{N}(0, 10)\ \forall p \in \boldsymbol{P}$$
$$\boldsymbol{\Sigma}_{intrinsic_{C_i}} \sim InverseWishart(\cdot)\ \forall C_i \in \boldsymbol{C}$$
$$\boldsymbol{\Sigma}_{extrinsic} \sim InverseWishart(\cdot)$$
$$\boldsymbol{\mu}_p \sim \mathcal{N}(\boldsymbol{\phi}_p, \boldsymbol{\Sigma}_{extrinsic})$$
$$\boldsymbol{y}_{n_{C_i}} \sim \mathcal{N}(\boldsymbol{\mu}_{x_{n_{C_i}}}, \boldsymbol{\Sigma}_{intrinsic_{C_i}})$$

Here,

$$\boldsymbol{\Phi} = \begin{bmatrix} \boldsymbol{\phi}_1 \\ \boldsymbol{\phi}_2 \\ \vdots \\ \boldsymbol{\phi}_P \end{bmatrix} \quad \boldsymbol{M} = \begin{bmatrix} \boldsymbol{\mu}_1 \\ \boldsymbol{\mu}_2 \\ \vdots \\ \boldsymbol{\mu}_P \end{bmatrix} \quad \boldsymbol{Y}_{C_i} = \begin{bmatrix} \boldsymbol{y}_1 \\ \boldsymbol{y}_2 \\ \vdots \\ \boldsymbol{y}_{n_{C_i}} \end{bmatrix}$$

<u>Target function</u>

The target function of the model:

$$\mathbb{P}(\boldsymbol{\Phi}, \boldsymbol{M} \mid \boldsymbol{Y}) \propto \mathbb{P}(\boldsymbol{W}, \boldsymbol{H}, \boldsymbol{M}, \boldsymbol{\Sigma} \mid \boldsymbol{Y}) \propto \mathbb{P}(\boldsymbol{Y} \mid \boldsymbol{M}, \boldsymbol{\Sigma})\mathbb{P}(\boldsymbol{M} \mid \boldsymbol{W}, \boldsymbol{H})\mathbb{P}(\boldsymbol{W})\mathbb{P}(\boldsymbol{H})\mathbb{P}(\boldsymbol{\Sigma})$$

The joint posterior is then:

$$\mathbb{P}(\boldsymbol{\Phi}, \boldsymbol{M} \mid \boldsymbol{Y}) \propto \left(\prod_{i=1}^{C} \mathbb{P}(\boldsymbol{\Sigma_i})\mathbb{P}(\boldsymbol{Y_i} \mid \boldsymbol{M}, \boldsymbol{\Sigma_i})\right) \cdot \left(\prod_{i=1}^{P} \mathbb{P}(\boldsymbol{M} \mid \boldsymbol{W}, \boldsymbol{H_i}) \cdot \mathbb{P}(\boldsymbol{H_i})\right) \cdot \left(\prod_{i=1}^{L} \mathbb{P}(\boldsymbol{W_i})\right) \propto$$
$$\left(\prod_{i=1}^{C} \prod_{n=1}^{N_i} \mathbb{P}(\boldsymbol{\Sigma_i})\mathbb{P}(\boldsymbol{Y}_n \mid \boldsymbol{M}_{x_n}, \boldsymbol{\Sigma_i})\right) \cdot \left(\prod_{i=1}^{P} \mathbb{P}(\boldsymbol{M} \mid \boldsymbol{W}, \boldsymbol{H_i}) \cdot \mathbb{P}(\boldsymbol{H_i})\right) \cdot \left(\prod_{i=1}^{L} \mathbb{P}(\boldsymbol{W_i})\right)$$

To encourage the model to find more distinct niches, we add an orthogonality term to the model target function. The upper triangular matrix of the transpose cross product of the niche basis matrix $\boldsymbol{W}$ is used to calculate a mean as a proxy for niche similarity. This term is weighted by a hyperparameter $\lambda$ and minimized in the target function. Therefore, using maximum a posteriori inference, we maximize the new target function including the log joint posterior and the orthogonality term:

$$\sum_{i=1}^{C} \left( log\mathbb{P}(\boldsymbol{\Sigma_i}) + \sum_{n=1}^{N_i} log\mathbb{P}(\boldsymbol{Y}_n \mid \boldsymbol{M}_{x_n}, \boldsymbol{\Sigma_i}) \right) + \sum_{i=1}^{P} (log\mathbb{P}(\boldsymbol{M} \mid \boldsymbol{W}, \boldsymbol{H_i}) + log\mathbb{P}(\boldsymbol{H_i}))$$
$$+ \sum_{i=1}^{L} log\mathbb{P}(\boldsymbol{W_i}) + \lambda \cdot Mean\big(upper\_tri(\boldsymbol{W}^{\intercal}\boldsymbol{W})\big)$$

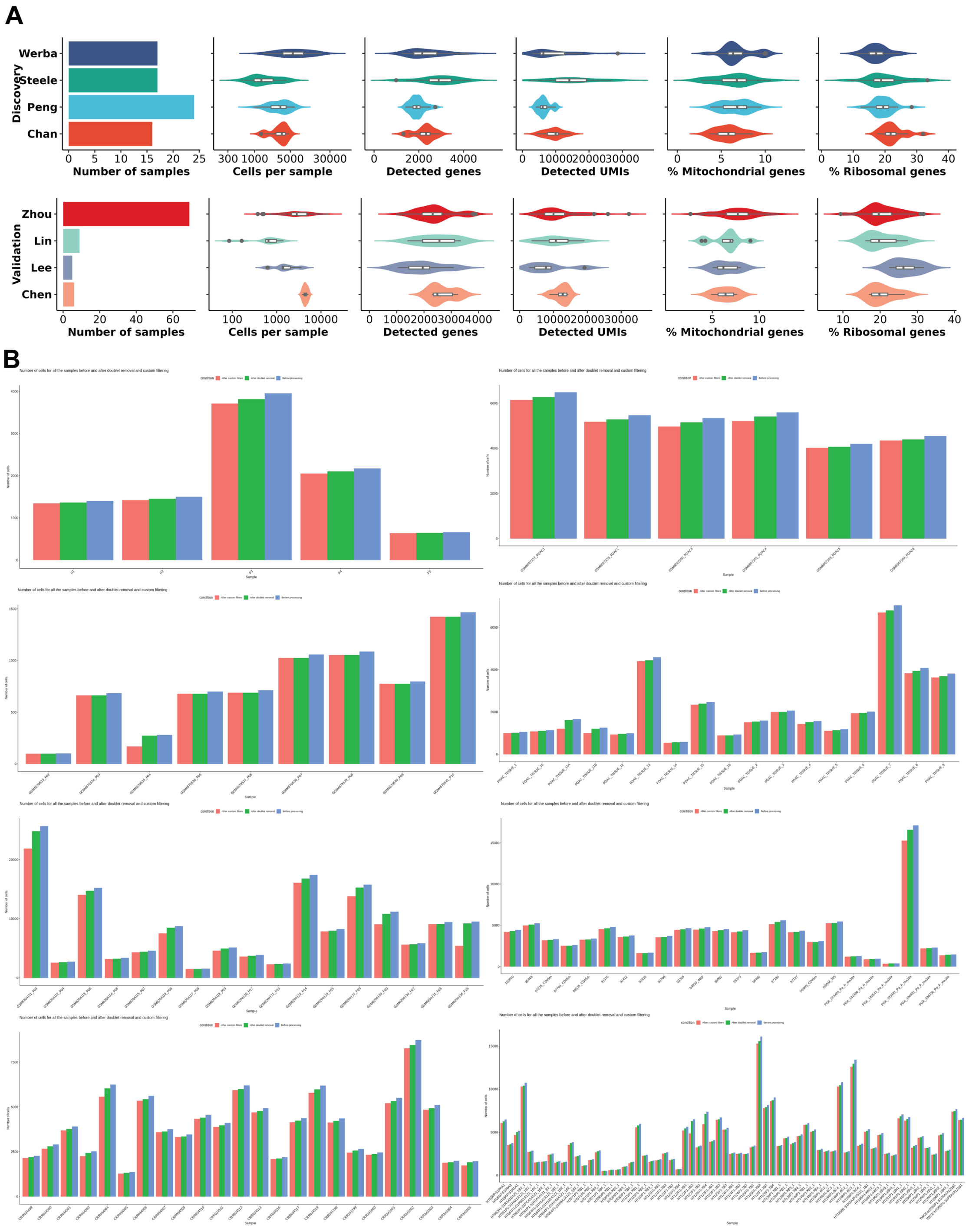


**Fig. S1.**

**A**, Dataset and quality control summary for the single-cell PDAC atlas. **B**, Bar plots showing the number of cells in each primary tumour sample before quality control (blue), after doublet removal (green), and after doublet removal and quality filtering (red). Each panel shows samples in one patient cohort.

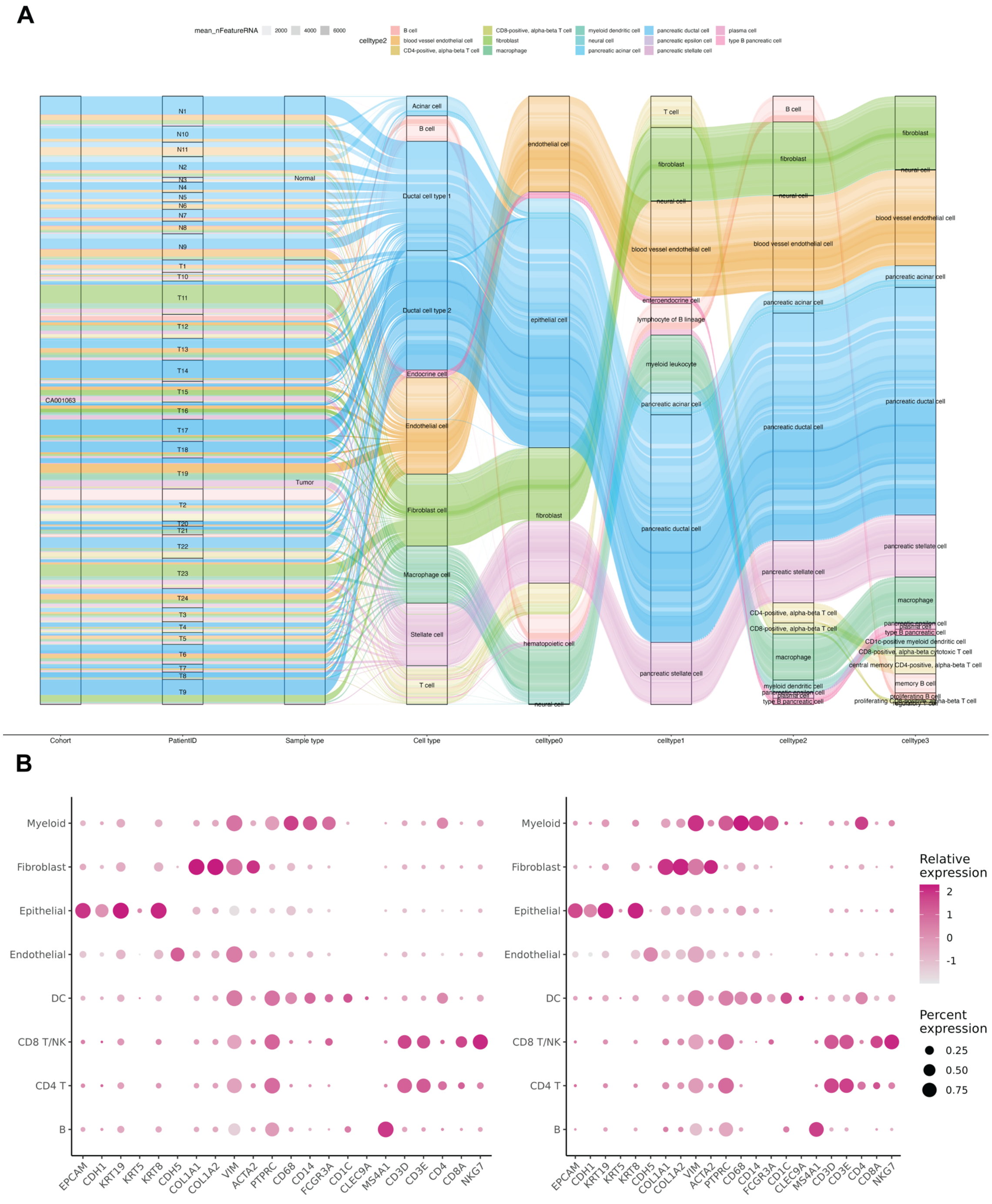


**Fig. S2.**

**A**, Cell type information fields in the PDAC single cell atlas used as reference for in house annotation. **B**, Cell type marker expression in consolidated cell types in discovery datasets (left) and validation datasets (right).

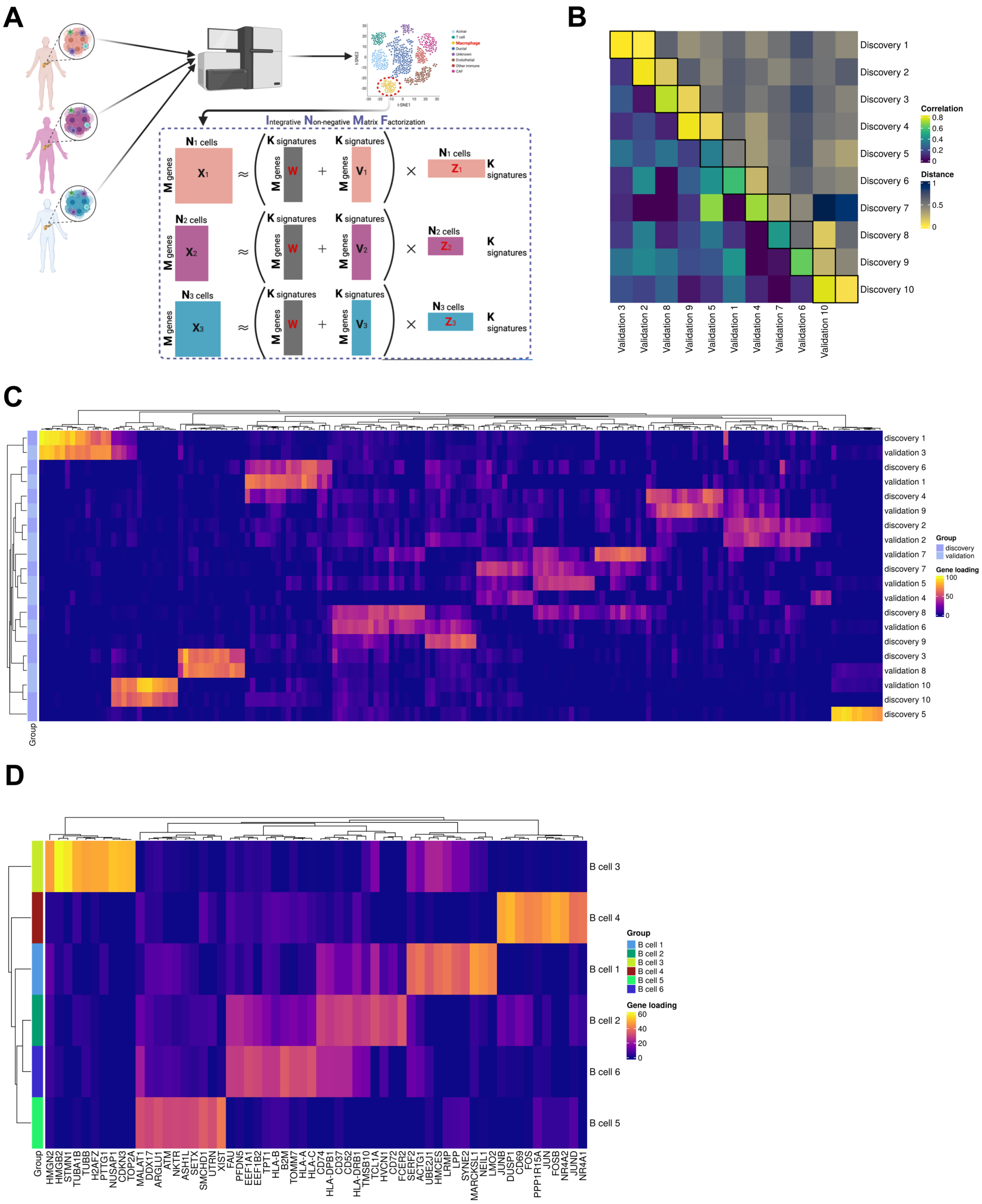


**Fig. S3.**

**A**, Schematic for cell type-specific gene programs extraction by iNMF. **B**, Gene weights correlations and Manhattan distances between candidate fibroblast gene programs extracted from the discovery datasets and the validation datasets. Columns (validation gene programs) are ordered according to best matching discovery-validation gene program pairs. **C**, Top 10 gene loadings in the fibroblast programs obtained from fibroblasts in the discovery datasets (discovery 1 to discovery 10) and the fibroblast programs obtained from the fibroblasts in the validation datasets (validation 1 to validation 10). **D**, Top 10 loaded genes in the B cell programs after removing programs with no match in the validation datasets.

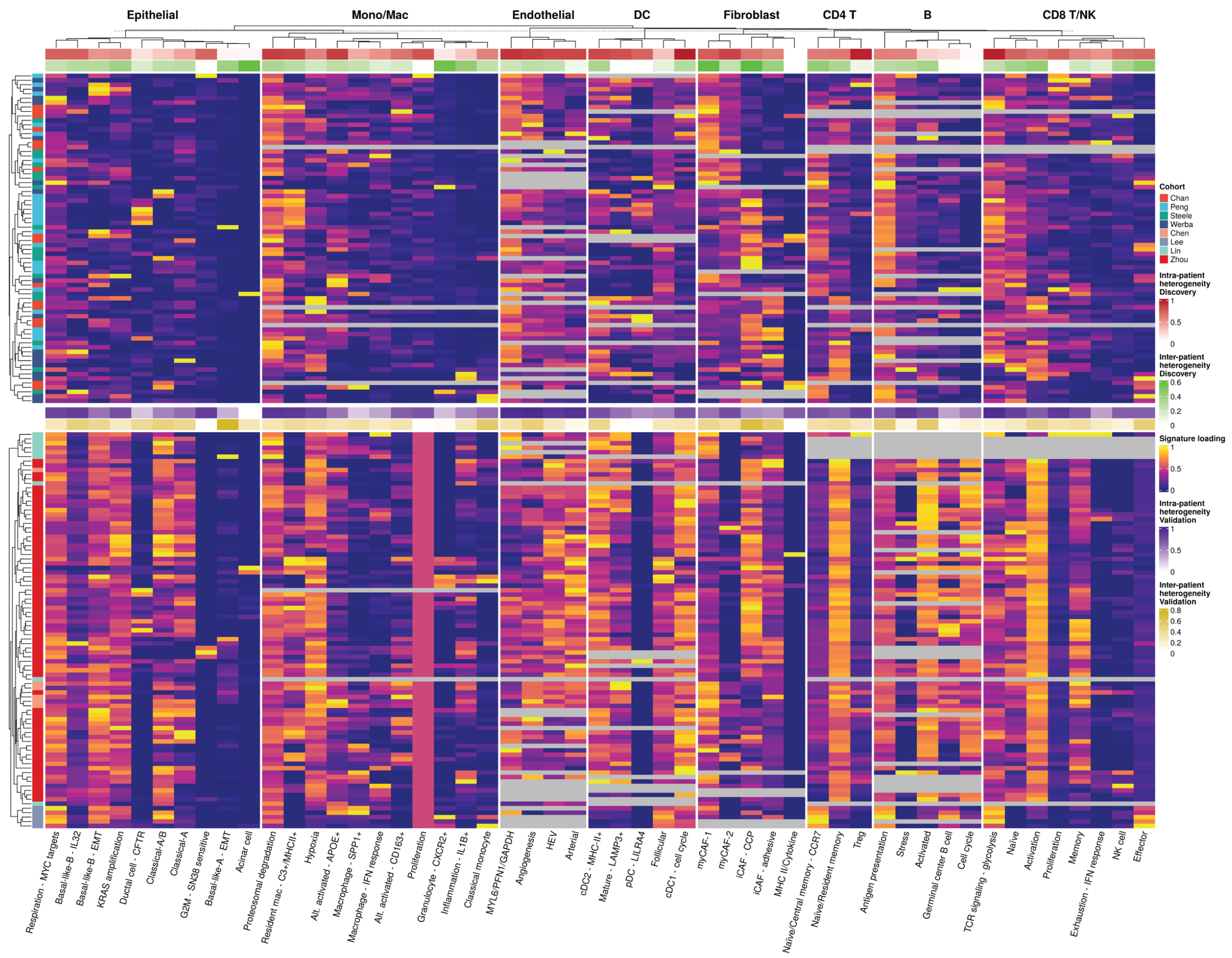


**Fig. S4.**

Consolidated gene program sample-level loading weights. Each row is a different tumour sample; the left bar indicates patient cohort. The samples are grouped by discovery/validation. Each column is a cell type specific gene program labeled according to contextualization results; the top bar indicates sample-level gene program loading variance means in discovery; the second bar indicates gene program loading mean variance between samples in discovery (**Methods**). The heatmap denotes scaled loadings of each gene program for each tumour sample, grouped by cell types indicated on the top.

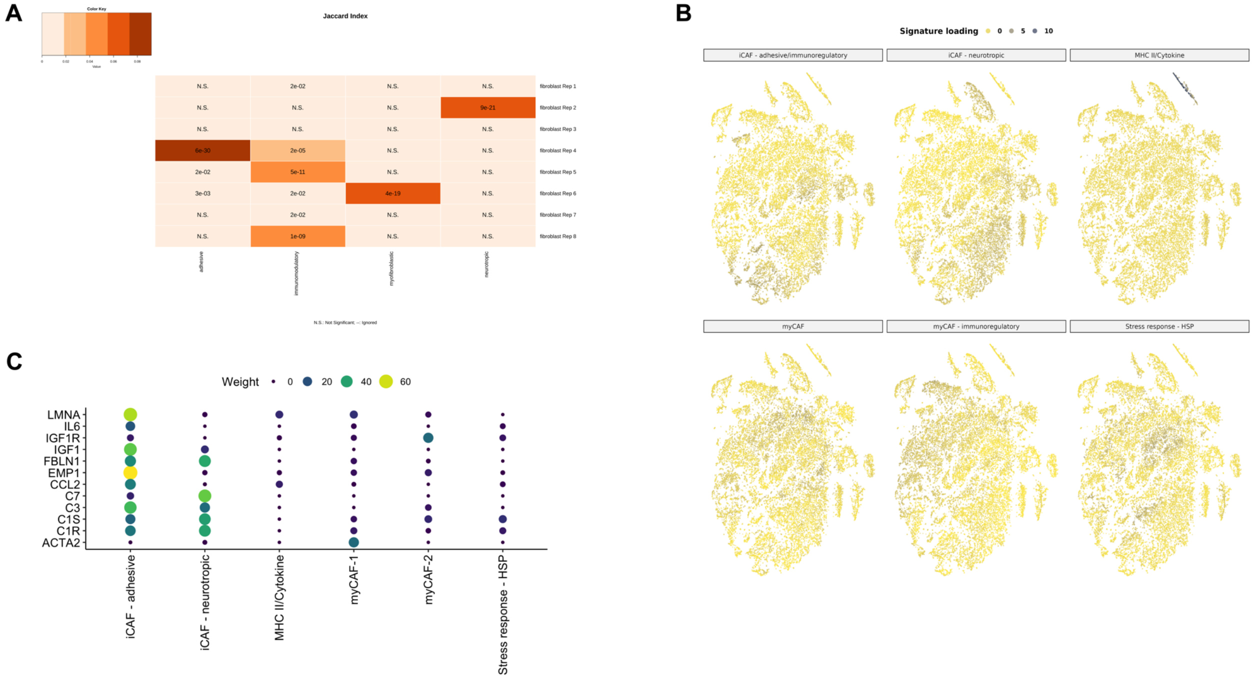


**Fig. S5.**

**A**, Gene set overlap analysis between the top 50 loaded genes of fibroblast gene programs and published PDAC fibroblast cell state marker gene lists. **B**, Raw loadings of fibroblast gene programs in single cells in the discovery cohorts. **C**, Selected gene weights in fibroblast gene programs.

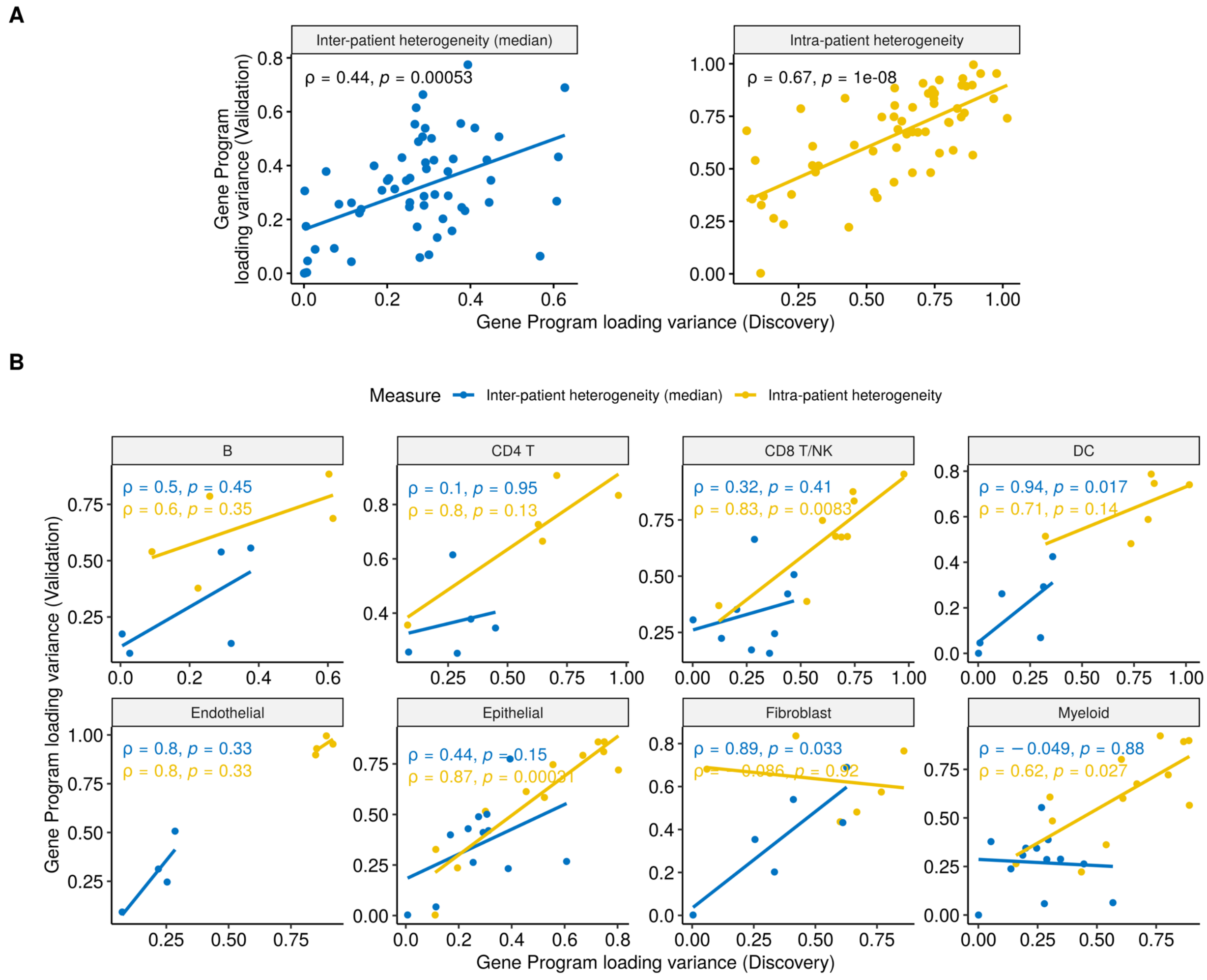


**Fig. S6.**

**A**, Spearman correlations of overall gene program loading heterogeneity. Each dot is one cell type specific program. Inter-patient heterogeneity (left) was calculated by first getting the program loading means in each sample, then computing the variance of sample level loading means in discovery and validation, respectively. Intra-patient heterogeneity (right) was calculated by first computing the program loading variances in each sample, then getting the mean of sample level loading variances in discovery and validation, respectively. **B**, Spearman correlations of gene program loading heterogeneity grouped by cell lineage. Each dot is one gene program.

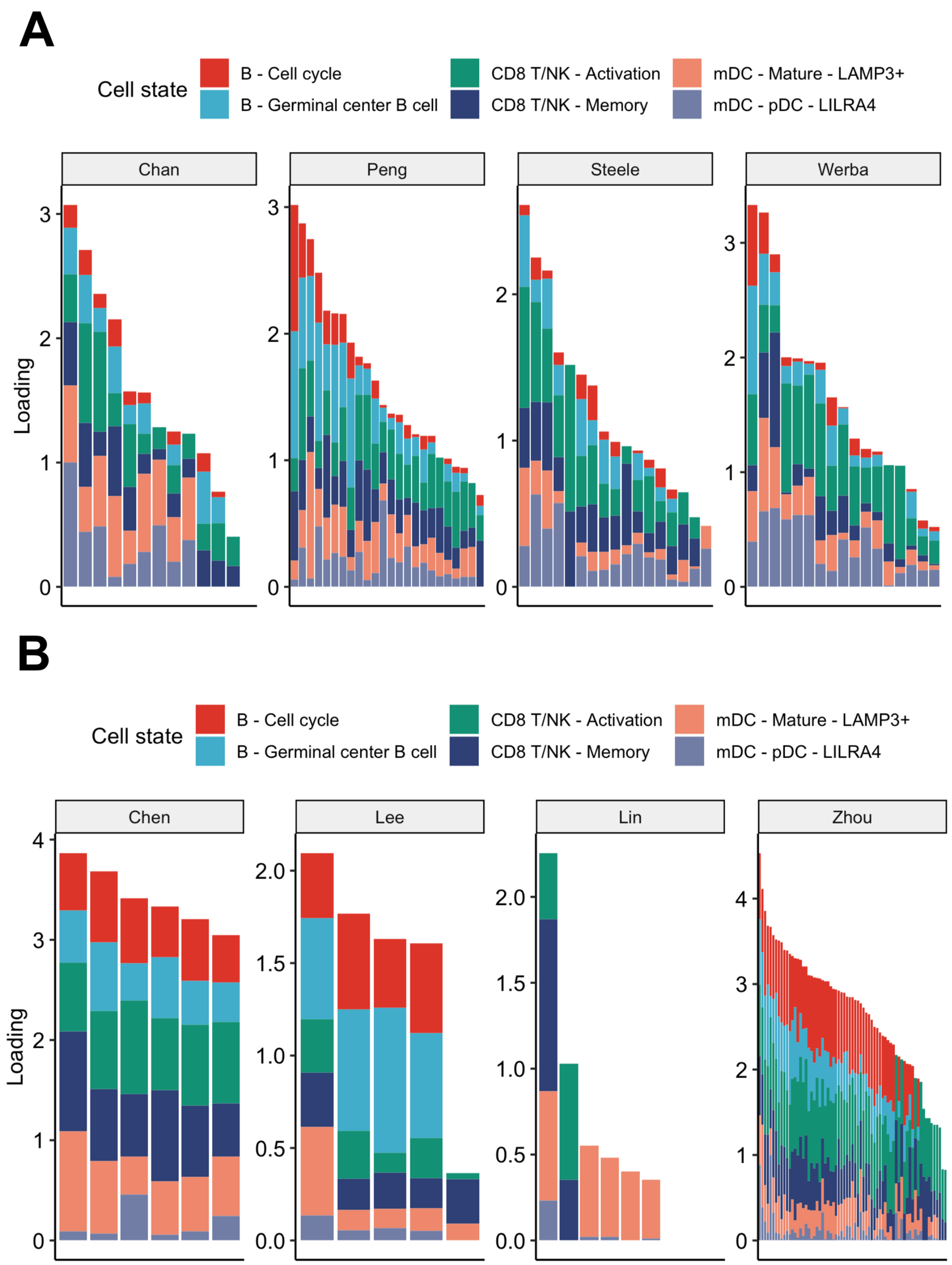


**Fig. S7.**

**A and B**, Loadings of the TLS related programs in the discovery patient samples (**A**) and in the validation patient samples (**B**).

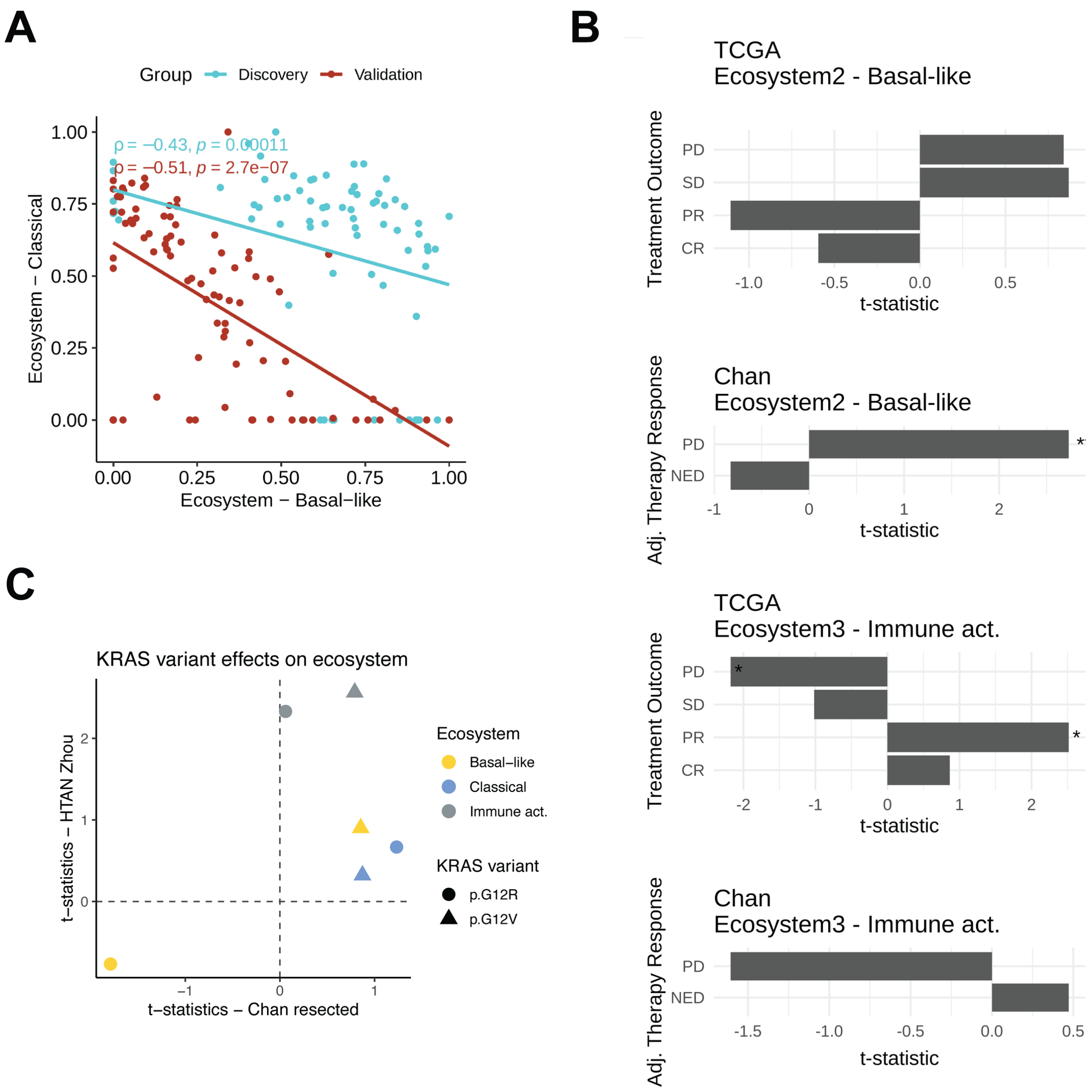


**Fig. S8.**

**A**, Loadings of Ecosystems in scRNA PDAC tumour samples. **B**, T-statistics from linear regression models fitting relationships between treatment outcome and ecosystem loadings in bulk RNA-seq tumour samples. **C**, T-statistics from linear regression models fitting relationships between KRAS variants and ecosystem loadings in bulk RNA-seq tumour samples.

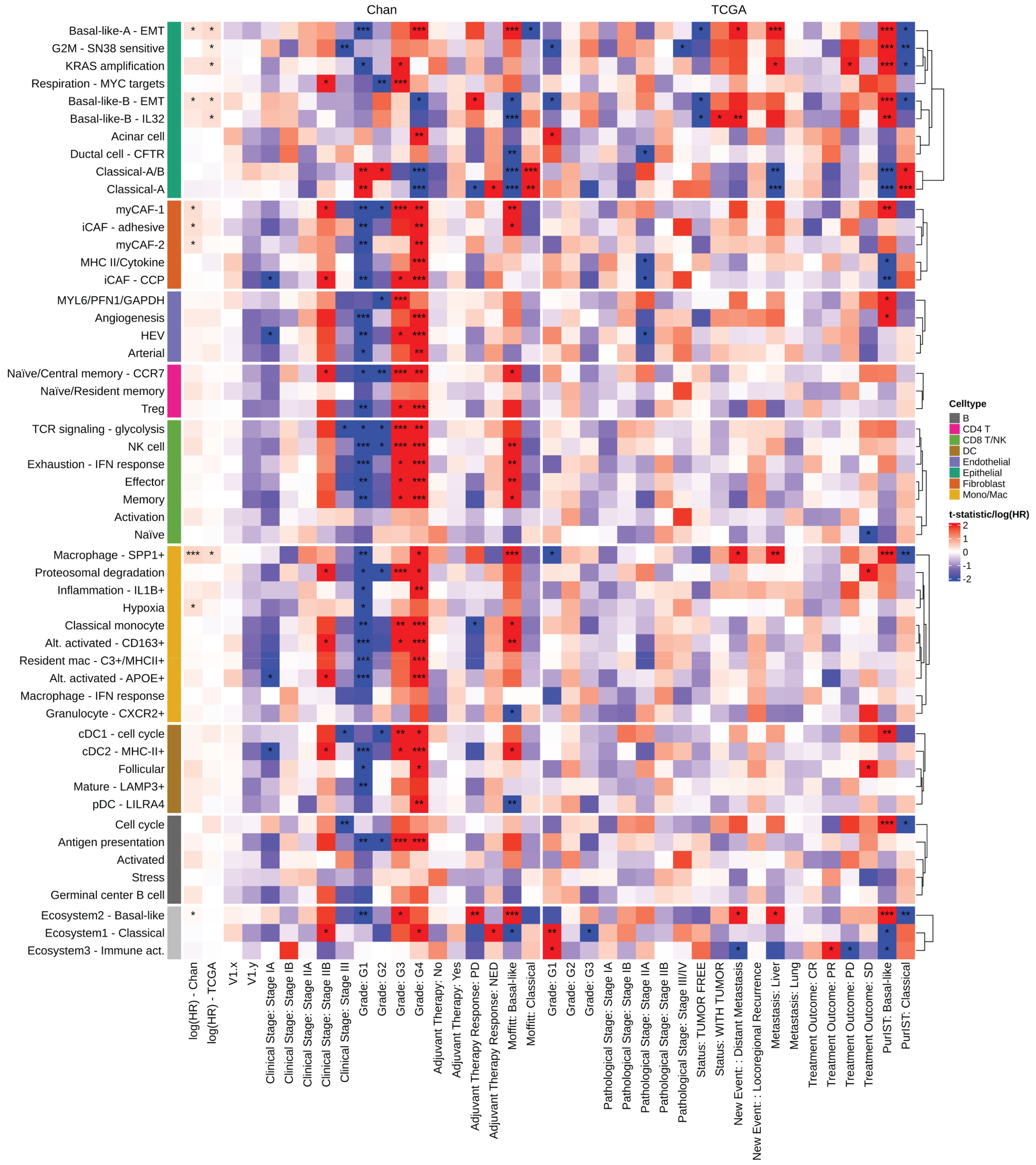


**Fig. S9.**

Clinical associations of cell type-specific gene programs and multi-cellular ecosystems. The two leftmost columns of the heatmap show log(HR) in the bulk RNA-seq cohorts from fitting univariate CoxPH models. The other columns of the heatmap show fitted coefficients for each program and ecosystems using linear regression. '*': p-value < 0.05; '**': p-value < 0.01, '***': p-value < 0.001 (all Benjamini-Hochberg FDR corrected).

**Table S1. (separate file)**
Basic information on the eight curated scRNA-seq datasets.

**Table S2. (separate file)**
Weights of genes in reproducible cell type specific programs.

**Table S3. (separate file)**
Linear model estimates of KRAS variant effect sizes on multi-cellular niche loadings in patient samples.

**Table S4. (separate file)**
Clinical associations of all cell state programs.
